# NAD^+^ Sensing by PARP7 Regulates the C/EBPβ-Dependent Transcription Program in Adipose Tissue In Vivo

**DOI:** 10.1101/2025.04.07.647692

**Authors:** MiKayla S. Stokes, Yoon Jung Kim, Yonghyeon Kim, Sneh Koul, Shu-Ping Chiu, Morgan Dasovich, Josue Zuniga, Tulip Nandu, Dan Huang, Thomas P. Mathews, Ashley Solmonson, Cristel V. Camacho, W. Lee Kraus

## Abstract

We have identified PARP7, an NAD^+^-dependent mono(ADP-ribosyl) transferase, as a key regulator of the C/EBPβ-dependent proadipogenic transcription program. Moreover, PARP7 is required for efficient adipogenesis and downstream biological functions, including involution of the lactating mammary gland. PARP7 serves as a coregulator of C/EBPβ, and depletion of PARP7 causes a dramatic reduction in C/EBPβ binding across the genome. PARP7 functions as a sensor of nuclear NAD^+^ levels to control gene expression. At the relatively high nuclear NAD^+^ concentrations in undifferentiated preadipocytes, PARP7 is catalytically active for auto-mono(ADP-ribosyl)ation (autoMARylation). As nuclear NAD^+^ concentrations decline post-differentiation, autoMARylation decreases dramatically. AutoMARylation promotes instability of PARP7 through an E3 ligase-ubiquitin-proteasome pathway mediated by the ADP-ribose (ADPR)-binding ubiquitin E3 ligases DTX2 and RNF114. Genetic depletion of PARP7 in mice promotes a dramatic reduction in a wide array of lipids in the mammary gland fat pads and milk from lactating females, as well as a significant decrease in nicotinamide mononucleotide (NMN), a key nutrient in mother’s milk. The latter is due to reduced expression of *Nampt*, the gene encoding NAMPT, the enzyme that produces NMN, which is a direct transcriptional target of PARP7 and C/EBPβ. Collectively, our results extend the biology of PARP7 to adipogenesis and perinatal health. Moreover, our results describe the molecular events that regulate these downstream biological functions.

## Introduction

Adipogenesis, the cellular process by which a preadipocyte differentiates into a mature fat-storing adipocyte, is an important component of adipose tissue health (Ghaben and Scherer, 2019). Dysregulation of adipogenesis and adipose tissue homeostasis is an underlying cause of human diseases, including metabolic syndrome and obesity, as well as cancer and reproductive issues (Brown and Scherer, 2023; Dumesic et al., 2016). Greater understanding of the metabolic pathways, signaling cascades, and molecular mechanisms that control adipogenesis has the potential to drive the development of new therapeutic interventions and improve human health. While key facets of the molecular and cellular mechanisms underlying adipogenesis have been elucidated, other aspects have not been well characterized (Ambele et al., 2020). Although ADP-ribosylation (ADPRylation) and the PARP family of enzymes that mediate this modification have been implicated in adipogenesis (Szanto and Bai, 2020; Szanto et al., 2021), many details remain poorly characterized.

ADPRylation is a posttranslational modification (PTM) of proteins that results in the covalent linkage of ADP-ribose (ADPR) moieties from oxidized β-nicotinamide mononucleotide (NAD^+^) on a variety of residues (e.g., Glu, Asp, Ser) on substrate proteins (Cohen and Chang, 2018; Gupte et al., 2017). ADPRylation may occur via the attachment of a single ADPR moiety [i.e., monoADPRylation (MARylation) mediated by mono(ADP-ribosyl) transferases or MARTs] or multiple ADPR moieties [i.e., oligoADPRylation or polyADPRylation (PARylation) mediated by poly(ADP-ribosyl) polymerases or PARPs] (Gibson and Kraus, 2012). Since ADPRylation consumes NAD^+^, the resynthesis of NAD^+^ is essential to maintain the activity of PARP enzymes. Importantly, NAD^+^ biosynthesis is highly compartmentalized in the cell, with distinct nuclear and cytoplasmic pools generated by distinct nuclear and cytoplasmic nicotinamide adenylyl transferases (i.e., NMNAT-1 and NMNAT-2, respectively) (Cambronne and Kraus, 2020).

We and others have previously linked nuclear ADPRylation by PARP1 to metabolic phenotypes and adipogenesis in cultured cells and mice (Szanto and Bai, 2020; Szanto et al., 2021). We have also previously defined a definitive set of nuclear substrate proteins whose ADPRylation by PARP1 regulates the differentiation of preadipocytes to mature adipocytes. For example, PARP1-mediated ADPRylation of C/EBPβ, a key proadipogenic transcription factor, inhibits its transcriptional activity and prevents the differentiation of preadipocytes (Luo et al., 2017; Ryu et al., 2018). Likewise, PARP1-mediated ADPRylation of histone H2B at Glu35, which inhibits AMPK-mediated phosphorylation on H2B at Ser36, inhibits the proadipogenic transcriptional program and prevents the differentiation of preadipocytes (Huang et al., 2020). Rapidly declining nuclear NAD^+^ levels in response to differentiation signals attenuates PARP1 activity and releases these ADPRylation-mediated inhibitory mechanisms, allowing for differentiation to proceed (Ryu et al., 2018). The role of other nuclear PARPs and MARylation in the regulation of proadipogenic gene expression has not been well characterized.

PARP7 (a.k.a. 2,3,7,8-tetrachlorodibenzodioxin-inducible PARP or TIPARP), a MART with nuclear functions in gene regulation, has recently received increased attention since the development of high affinity, high selectivity inhibitors, such as RBN2397 (Gozgit et al., 2021) and KMR-206 (Sanderson et al., 2023). The gene regulatory actions of PARP7 have been studied in the context of the aryl hydrocarbon receptor (AHR) (Ahmed et al., 2015; MacPherson et al., 2013), estrogen receptor alpha (ERα) (Rasmussen et al., 2021), and androgen receptor (AR) (Kamata et al., 2021b; Yang et al., 2021). AR is MARylated by PARP7, and the MARylated sites are bound by the ADPR-reading macrodomains in PARP9, which complexes with DTX3L, a Deltex E3 ubiquitin ligase (Yang et al., 2021). Thus, PARP7 acts as an enzyme, a coregulator, and a scaffold to bring together multiple transcriptional regulatory proteins in a complex to control signal-regulated gene expression. Biologically, PARP7 is a stress-responsive regulatory protein that has been implicated in stem cell pluripotency, viral replication, neuronal function, the regulation of innate and adaptive immunity, and oncology (Cohen and Chang, 2018; Grimaldi et al., 2019; Jeltema et al., 2025; Manetsch et al., 2023; Roper et al., 2014; Yamada et al., 2016). Here, we identify PARP7 as a key regulator of the C/EBPβ-dependent proadipogenic transcription program that is required for efficient adipogenesis and downstream biological functions, including involution of the lactating mammary gland.

## Results

### PARP7 is required for adipogenesis

We have previously shown that NAD^+^-dependent, PARP1-mediated, site-specific ADPRylation of key nuclear substrate proteins (e.g., C/EBPβ, histone H2B) controls the differentiation of preadipocytes into mature adipocytes (Fig. 1A) (Huang et al., 2020; Luo et al., 2017; Ryu et al., 2018). Given the dramatic alterations in compartment-specific NAD^+^ synthesis and PARP1 activity that occur during adipogenesis (Ryu et al., 2018), we sought to determine if other members of the PARP family, specifically the MARTs, also regulate adipogenesis. To do so, we performed an siRNA screen to knockdown individual MARTs in 3T3-L1 cells, a preadipocyte cell line that can be differentiated into fat-accumulating mature adipocytes (Fig. 1A) (Green and Meuth, 1974). We examined if knockdown of individual MARTs (Suppl. Fig. S1A) altered adipogenesis based on a variety of commonly used endpoints, including the expression of two adipogenic marker genes, *Fabp4* and *Adipoq* (Suppl. Fig. S1B), as well as lipid accumulation monitored by Oil Red-O staining (Suppl. Fig. S1C). Although knockdown of several MARTs reduced adipogenesis based on these markers, knockdown of *Parp7* had the greatest effect (Fig. 1B; Suppl. Fig. S1). Thus, we focused our subsequent studies on PARP7, also known as TIPARP (Ma et al., 2001; MacPherson et al., 2013).

**Figure 1.**
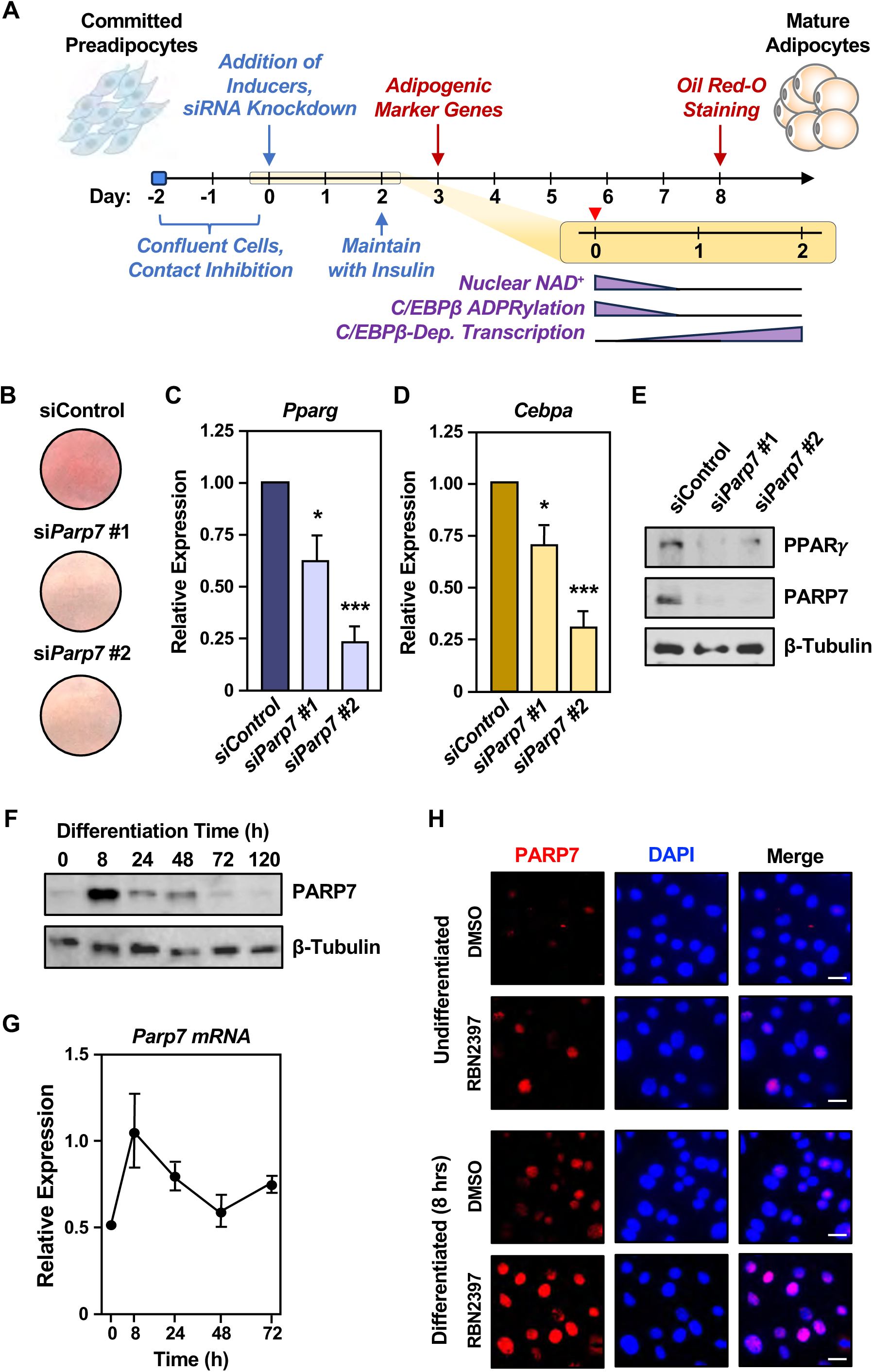
Depletion of PARP7 inhibits adipogenesis. **(A)** Schematic representation of the differentiation protocol used for 3T3-L1 cells. The timing of treatments, testing for biological endpoints, and key outcomes are indicated. **(B)** PARP7 is required for efficient differentiation and fat accumulation in 3T3-L1 cells. The cells were subjected to knockdown with control or *Parp7* siRNAs, followed by differentiation with MDI cocktail. Oil Red-O staining was performed to examine lipid accumulation on day 8 of differentiation. **(C and D)** PARP7 is required for efficient differentiation of 3T3-L1 cells. 3T3-L1 cells were subjected to knockdown with control or *Parp7* siRNAs, followed by differentiation with MDI cocktail. Bar graphs show expression of the mRNAs encoding adipogenic marker genes (C) *Pparg* and (D) *Cebpa* as assayed by RT-qPCR on day 3 of differentiation. Target gene expression was normalized to the expression of *Tbp* mRNA. Each bar represents the mean + SEM; n = 3. Bars marked with asterisks are significantly different from control; ANOVA; * = p < 0.0332, *** = p < 0.0002. **(E)** Western blot showing a reduction of PPARy and PARP7 protein levels at day 2 of differentiation in 3T3-L1 cells after knockdown with control or *Parp7* siRNAs. β-tubulin was used as a loading control. **(F)** Western blot showing PARP7 protein levels during a time course of 3T3-L1 cell differentiation. β-tubulin was used as a loading control. **(G)** Line graph showing the relative expression of *Parp7* mRNA during a time course of 3T3-L1 cell differentiation. Target gene expression was normalized to the expression of *Tbp* mRNA. Each point represents the mean ± SEM; n = 3. **(H)** Representative immunofluorescent images of undifferentiated and differentiated (8 hours) 3T3-L1 cells treated with DMSO or 400 nM RBN2397 showing PARP7 nuclear localization during 3T3-L1 cell differentiation. The DNA was stained with DAPI. Scale bar = 25 µm.

To confirm and expand these results, we performed additional experiments in 3T3-L1 cells and primary preadipocytes derived from the stromal vascular fraction (SVF) of white adipose tissue collected from mice (Kilroy et al., 2018). First, we confirmed the effects of *Parp7* knockdown in 3T3-L1 cells using two other marker genes, *Pparg* and *Cebpa* (Fig. 1, C and D), which encode well-known adipogenic regulators (Lefterova et al., 2008; Tontonoz et al., 1994). Second, we confirmed a reduction in PPARγ protein upon *Parp7* knockdown (Fig. 1E), as well as depletion of PARP7 protein using an affinity-purified homemade antibody to PARP7 (Fig. 1E; Suppl. Fig. S2). Third, we repeated key experiments in 3T3-L1 cells subjected to CRISPR/Cas9- mediated knockout of PARP7 and confirmed the inhibition of marker gene expression and lipid accumulation upon PARP7 depletion (Suppl. Fig. S3, A-D). Finally, we examined the effects of siRNA-mediated depletion of *Parp7* in SVF-derived mouse primary preadipocytes and confirmed the inhibition of marker gene expression and lipid accumulation upon PARP7 depletion (Suppl. Fig. S3, E-K). Interestingly, depletion of PARP7 did not affect the expression of *Cebpb,* but inhibited the expression of well-known genes that are regulated by C/EBPβ (e.g., *Cebpa*, *Pparg*, *Fabp4*, *Adipoq*). Collectively, these results indicate that PARP7 is required for the efficient differentiation of preadipocytes into mature adipocytes by supporting the C/EBPβ- mediated proadipogenic gene expression program.

### PARP7 levels are dynamic, and are regulated by NAD^+^ and ubiquitylation

We examined PARP7 expression and localization in 3T3-L1 cells and observed that the levels of PARP7 protein are maximal in the early phases (∼ 8 hours) of the differentiation process, returning to basal levels by 72 hours (Fig. 1F), a time when nuclear NAD^+^ levels are significantly reduced (Ryu et al., 2018). Although less dramatic, the levels of *Parp7* mRNA exhibited a similar pattern of expression (Fig. 1G). Immunofluorescent staining with our PARP7 antibody demonstrated nuclear localization of PARP7 with increased signal after 8 hours of differentiation, as well as treatment with the PARP7 inhibitor RBN2397 (Gozgit et al., 2021) (Fig. 1H). These results indicate that PARP7 (1) has a limited window of expression during adipogenesis, (2) is dynamically regulated, perhaps through protein stability, and (3) inhibition of its catalytic activity stabilizes the protein.

To explore these possibilities in more detail, we treated 3T3-L1 cells with the proteasome inhibitor MG132 with or without differentiation. As with RBN2397, we observed a dramatic stabilization of PARP7 in differentiated cells with MG132 treatment (Fig. 2A). Using the protein synthesis inhibitor cycloheximide (CHX), we determined a half-life of PARP7 of ∼8 minutes (Fig. 2, B and C), which is similar to a previous determination of ∼5 minutes (Kamata et al., 2021a). In PARP7 immunoprecipitation-Western blotting (IP-Western) experiments in 3T3- L1 cells, we observed that MG132-induced stability of PARP7 is associated with decreased autoMARylation of PARP7 (Fig. 2D). Moreover, treatment with nicotinamide mononucleotide (NMN), an NAD^+^ precursor that increases NAD^+^ levels in 3T3-L1 cells (Ryu et al., 2018) as indicated by enhanced autoPARylation of PARP1 (Fig. 2E, Input), increased PARP7 autoMARylation and decreased PARP7 protein levels (Fig. 2E, IP-PARP7). We determined the Km of PARP7 for NAD^+^ to be ∼70 µM using a biochemical assay (Fig. 2, F and G), a value that is between the high (∼100 µM, undifferentiated) and low (∼40 µM, 8 hours of differentiation) concentrations of nuclear NAD^+^ in 3T3-L1 cells that we determined previously (Ryu et al., 2018). Together, these results indicate that PARP7 autoMARylation, driven by changes in NAD^+^ concentrations, is associated with decreased PARP7 stability. These results are consistent with previous studies showing that catalytically dead PARP7 is expressed at a much higher level compared to wild-type (WT) PARP7 (Gomez et al., 2018) and that chemical inhibition of PARP7 catalytic activity dramatically stabilizes PARP7 protein levels (Sanderson et al., 2023).

**Figure 2.**
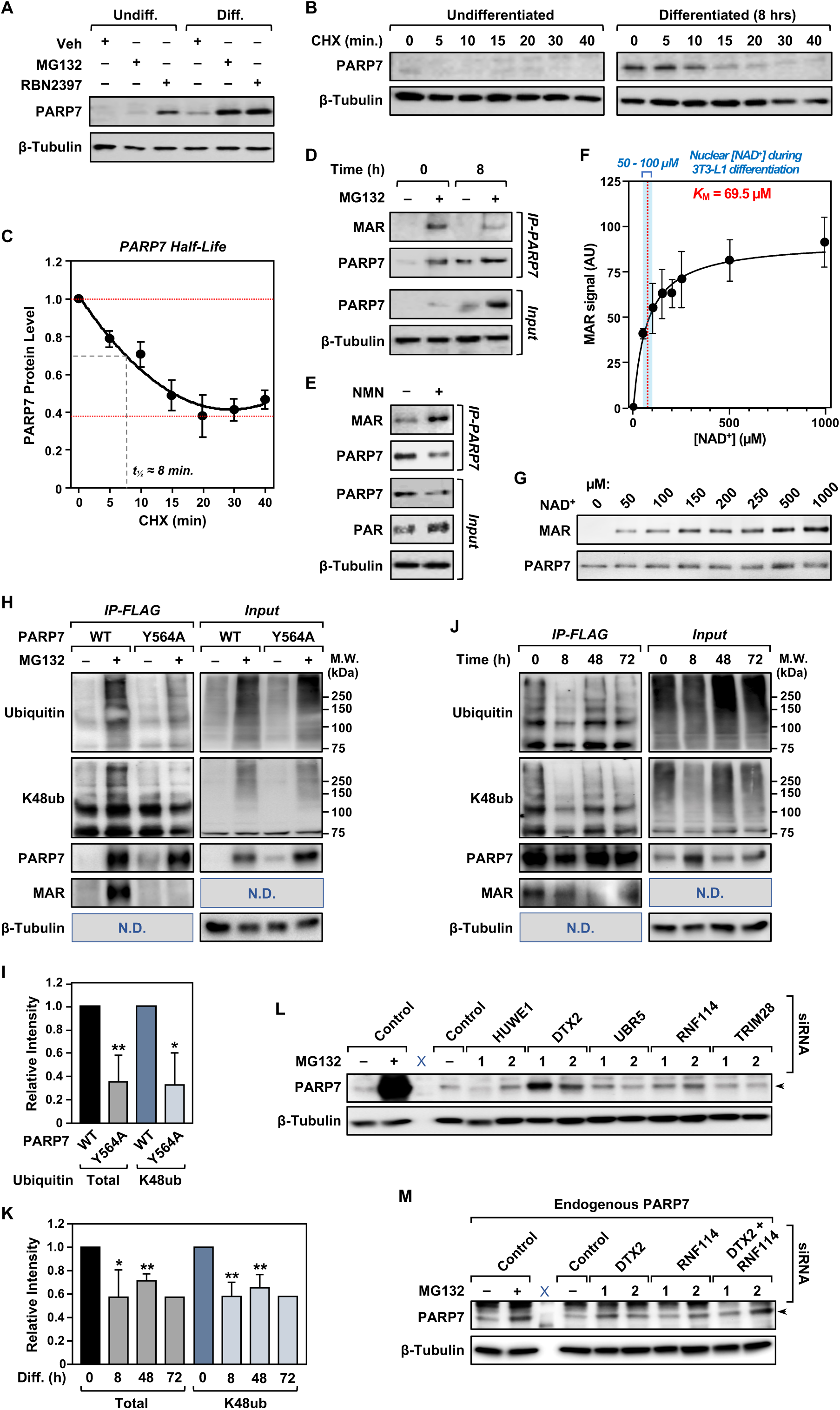
PARP7 protein stability is regulated by NAD^+^ availability, autoMARylation, ubiquitylation, and proteasome-mediated degradation. **(A)** PARP7 is stabilized by inhibition of the proteasome and PARP7 catalytic activity. Differentiation of 3T3-L1 cells was initiated using MDI cocktail and the cells were subjected to treatment with DMSO vehicle, 10 µM MG132, or 400 nM RBN2397 for 8 hours. The cells were collected after 8 hours of differentiation or treatment without differentiation. Western blot showing PARP7 protein levels. β-tubulin was used as a loading control. **(B and C)** Time course of PARP7 stability. 3T3-L1 cells were differentiated using MDI cocktail. At 0 or 8 hours of differentiation, the cells were subjected to treatment with 100 μg/mL cycloheximide (CHX) for the lengths of time indicated and collected. (B) Western blotting for PARP7 at 0 or 8 hours of differentiation revealed the reduction of PARP7 protein levels in the absence of ongoing protein synthesis. (C) Line graph showing the quantification of PARP7 Western blots like those shown in panel (B) at 8 hours of differentiation. Each point represents the mean ± SEM; n = 3. **(D and E)** Enhanced PARP7 autoMARylation and NAD^+^ sensing destabilizes PARP7. 3T3-L1 cells with Dox-inducible ectopic expression of PARP7 were subjected to treatment with (D) DMSO vehicle or 10 µM MG132 or (E) 10 mM NMN after differentiation using MDI cocktail. At 0 or 8 hours of differentiation, the cells were collected and PARP7 was immunoprecipitated. Undifferentiated samples received the same 8 hour treatments of vehicle or 10 µM MG132. Western blots showing PARP7 automodification (MAR), PARP7, and PAR. β-tubulin was used as a loading control. **(F and G)** The K_m_ of PARP7 for NAD^+^ is near the midpoint of high and low nuclear NAD^+^ concentrations in differentiating adipocytes as determined previously (Ryu et al., 2018). (F) Graph showing a K_m_ determination using purified recombinant PARP7 in a biochemical assay with increasing amounts of NAD^+^ and Western blotting for MAR like the assay shown in (G). In (F), each point represents the mean ± SEM; n = 5. (**H and I)** IP-Western assays showing PARP7 MARylation and ubiquitylation (total and K48- linked) with or without proteasome inhibition. (H) Undifferentiated 3T3-L1 cells with Dox- inducible ectopic expression of wild-type (WT) or catalytically dead mutant (Y564A) FLAG- tagged PARP7 were treated ± 10 µM MG132 for 6 hours. PARP7 was immunoprecipitated and subjected to Western blotting for MAR and ubiquitylation, as indicated. β-tubulin was used as a loading control. (I) Quantification of multiple replicates like the one shown in panel (H). Each bar represents the mean + SEM; n = 3. Bars marked with asterisks are significantly different from control; Student’s t-test; * = p < 0.0122, ** = p < 0.007. **(J and K)** IP-Western assays showing PARP7 MARylation and ubiquitylation (total and K48- linked) during a time course of differentiation. (J) 3T3-L1 cells ectopically expressing FLAG- tagged WT PARP7 were subjected to Dox induction during a time course of differentiation as indicated. The cells were treated with 10 µM MG132 for 6 hours to stabilize PARP7. PARP7 was immunoprecipitated (IP-FLAG) and subjected to Western blotting for MAR and ubiquitylation, as indicated. β-tubulin was used as a loading control. (K) Quantification of multiple replicates like the one shown in panel (J). Each bar represents the mean + SEM; n = 3, except for 72 hours, where n = 1. Bars marked with asterisks are significantly different from control; Student’s t-test; * = p ≤ 0.03, ** = p ≤ 0.004. **(L)** A small-scale siRNA screen of ubiquitin E3 ligases identified DTX2 and RNF114 as E3 ligases for the stabilization of PARP7. Undifferentiated 3T3-L1 cells with Dox-induced ectopic expression of WT PARP7 were subjected to siRNA-mediated knockdown of one of five different E3 ubiquitin ligases. Western blot showing ectopically expressed PARP7 levels in cells with the indicated knockdown. siControl cells were treated ± 10 µM MG132 for 6 hours as indicated. β- tubulin was used as a loading control. **(M)** Western blot assays of endogenous PARP7. Undifferentiated 3T3-L1 were subjected to siRNA-mediated knockdown of *Dtx2*, *Rnf114*, or both. Western blot showing endogenous PARP7 protein levels. siControl cells were treated ± 10 µM MG132 as indicated. β-tubulin was used as a loading control.

Previous studies have shown that WT, but not catalytically dead, PARP7 is ubiquitylated, suggesting that the catalytic activity of PARP7 is necessary for autoMARylation-mediated degradation (Zhang et al., 2020). To explore the potential roles of autoMARylation and ubiquitylation in PARP7 stability during adipogenesis, we performed PARP7 IP-Western experiments, blotting for total and lysine 48 (K48)-linked ubiquitin on 3T3-L1 cells ectopically expressing WT PARP7 or a catalytically dead mutant (Y564A) (MacPherson et al., 2013) (Fig. 3A) and treated ± MG132. We observed that MG132-stabilized WT PARP7 was more highly autoMARylated and ubiquitylated than the catalytically dead mutant PARP7 (Fig. 2, H and I). Moreover, we found that PARP7 ubiquitylation decreased from 0 to 8 hours of differentiation, corresponding to the peak of PARP7 stability (Fig. 2, J and K).

**Figure 3.**
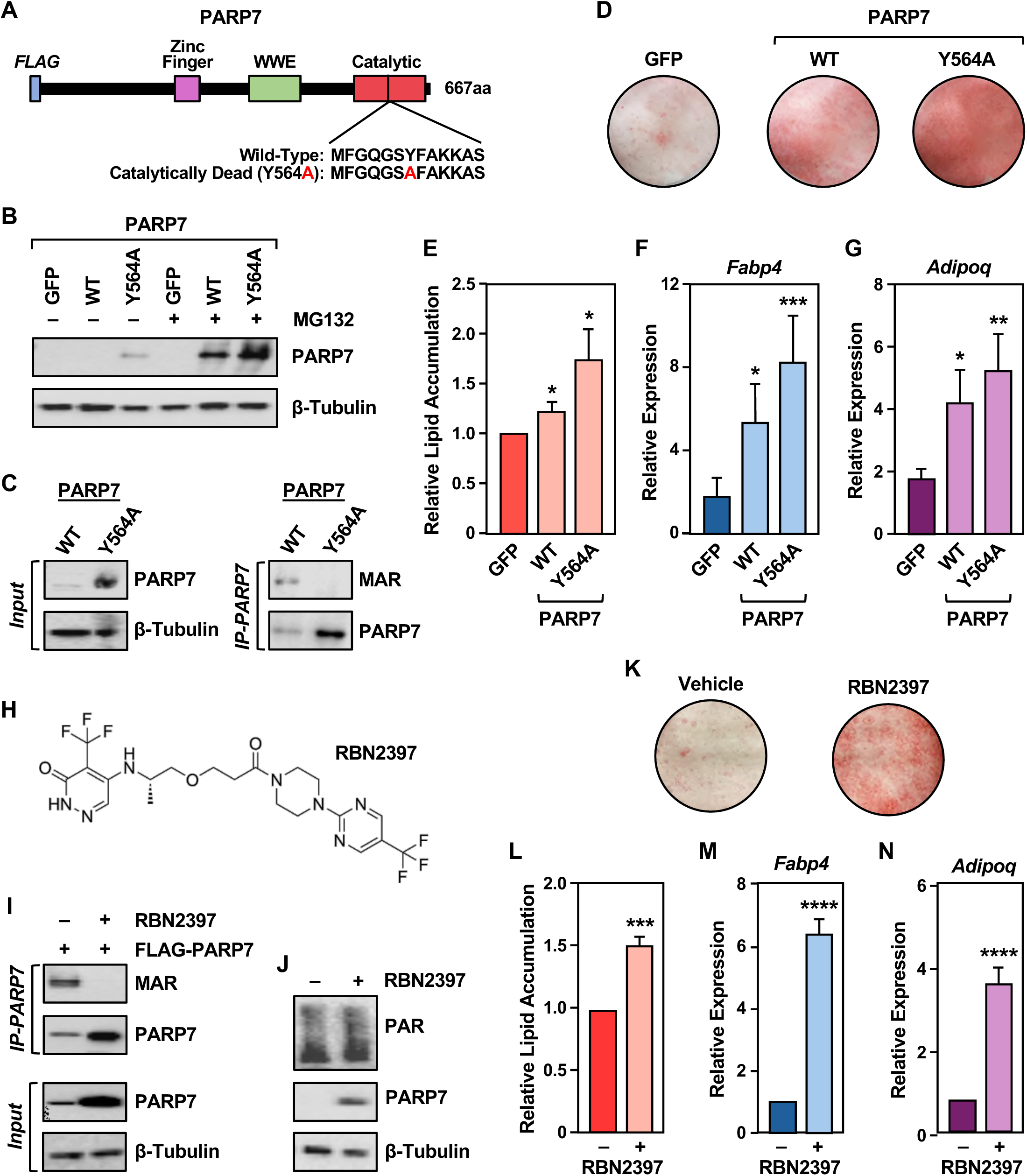
PARP7 catalytic activity is not required for adipogenesis. **(A)** Schematic representation of PARP7 showing the catalytic domain and the position of the Y564A point mutation that generates a catalytically dead version of PARP7. **(B)** Western blot showing PARP7 protein levels in undifferentiated 3T3-L1 ectopically expressing wild-type (WT) or catalytically dead mutant (Y564A) PARP7 upon Dox induction ± treatment with 10 µM MG132. β-tubulin was used as a loading control. **(C)** IP-Western assay showing MARylated PARP7 in HEK 293T cells with Dox-induced expression of WT or Y564A PARP7. β-tubulin was used as a loading control. **(D and E)** Catalytically dead PARP7 supports the differentiation of 3T3-L1 cells. (D) Lipid accumulation in 3T3-L1 cells with Dox-induced ectopic expression of WT or Y564A PARP7, or a GFP control, assayed by Oil Red-O staining at day 8 of adipocyte differentiation. (E) Quantification of multiple experiments like those shown in panel (D). Each bar represents the mean + SEM; n = 3. Bars marked with asterisks are significantly different from control; Student’s t-test; * = p < 0.0332. **(F and G)** Adipogenic marker gene expression in 3T3-L1 cells with Dox-induced ectopic expression of WT or Y564A PARP7, or a GFP control, after differentiation using MDI cocktail. Bar graphs showing mRNA expression of the adipogenic marker genes (F) *Fabp4* and (G) *Adipoq* assayed by RT-qPCR on day 3 of differentiation. Target gene expression was normalized to the expression of *Tbp* mRNA. Each bar represents the mean + SEM; n = 3. Bars marked with asterisks are significantly different from control; ANOVA; * = p < 0.0332, ** = p < 0.0021, *** = p < 0.0002. (H) Chemical structure of RBN2397, a PARP7 inhibitor. (I) IP-Western assay showing MARylated PARP7 in HEK 293T cells with Dox-induced expression of WT PARP7 ± treatment with 400 nM of RBN2397 for 16 hours. β-tubulin was used as a loading control. (J) Western blot showing PAR levels in 3T3-L1 cells treated with 400 nM RBN2397 for 8 hours at 8 hours of differentiation using MDI cocktail. β-tubulin was used as a loading control. **(K and L)** Treatment with RBN2397 enhances the differentiation of 3T3-L1 cells. (K) Lipid accumulation in 3T3-L1 cells treated with 400 nM RBN2397 throughout differentiation, assayed by Oil Red-O staining at day 8 of adipocyte differentiation. (L) Quantification of multiple experiments like those shown in panel (K). Each bar represents the mean + SEM; n = 3. Bars marked with asterisks are significantly different from control; Student’s t-test; *** = p < 0.0002. **(M and N)** Adipogenic marker gene expression in 3T3-L1 cells treated with 400 nM RBN2397 throughout differentiation using MDI cocktail. Bar graphs showing mRNA expression of the adipogenic marker genes (M) *Fabp4* and (N) *Adipoq* assayed by RT-qPCR on day 3 of differentiation. Target gene expression was normalized to the expression of *Tbp* mRNA. Each bar represents the mean + SEM; n = 3. Bars marked with asterisks are significantly different from control; ANOVA; **** = p < 0.0001.

We screened five E3 ubiquitin ligases previously shown to interact with PARP7 (i.e., HUWE1, DTX2, UBR5, RNF114, and TRIM28) (Zhang et al., 2020) using siRNA-mediated knockdown (Suppl. Fig. S3, L through R) in 3T3-L1 cells ectopically expressing PARP7. We observed that depletion of DTX2 and, to a lesser extent RNF114, increased the levels of PARP7 protein (Fig. 2L), indicating that they play a role in promoting the ubiquitin-mediated degradation of PARP7. Depletion of DTX2 or RNF114 also increased the levels of endogenous PARP7 protein in 3T3-L1 cells, with simultaneous depletion of both having an even greater effect (Fig. 2M). Collectively, these results indicate that PARP7 stability is regulated by ubiquitylation mediated by E3 ligases, including DTX2 and RNF114. Interestingly, both DTX2 and RNF114 contain domains known to interact with ADP-ribose (Li et al., 2023; Munzker et al., 2024).

### PARP7 catalytic activity is not required for adipogenesis

The results described above showed that PARP7 autoMARylation destabilizes PARP7, but did not address whether transmodification of substrate proteins by PARP7 is required for adipogenesis. PARP7 is known to MARylate various nuclear substrates in other biological systems, including histones (MacPherson et al., 2013) and transcription factors (e.g., liver X receptor, aryl hydrocarbon receptor, androgen receptor, and estrogen receptor) (Bindesboll et al., 2016b; Diani-Moore et al., 2010; Kamata et al., 2021b; Rasmussen et al., 2021). To determine if PARP7 catalytic activity is required for adipogenesis, we used a catalytically dead mutant of PARP7, Y564A (MacPherson et al., 2013) (Fig. 3A) and the PARP7-specific inhibitor, RBN2397 (Gozgit et al., 2021). When ectopically expressed in 3T3-L1 cells, the Y564A mutant exhibited greater stability than WT PARP7 (Fig. 3, B and C), as reported previously (Gomez et al., 2018), and an absence of autoMARylation, as expected (Fig. 3C). The increased stability of the Y564A mutant was further enhanced by treatment with MG132 (Fig. 3B). In spite of the loss of the catalytic activity, ectopic expression of the Y564A mutant promoted adipogenesis more than WT, as assessed by Oil Red-O staining (Fig. 3, D and E), and adipogenic marker gene expression, as assessed by RT-qPCR (Fig. 3, F and G). Similar results were observed with the PARP7 inhibitor RBN2397 (Fig. 3H). Namely, RBN2397 inhibited PARP7 autoMARylation and enhanced PARP7 stability (Fig. 3, I and J), while promoting adipogenesis, as assessed by Oil Red-O staining (Fig. 3, K and L), and adipogenic marker gene expression, as assessed by RT-qPCR (Fig. 3, M and N). Collectively, these results demonstrate that PARP7 catalytic activity is not required for the promotion of adipogenesis. Rather, they indicate that inhibition of PARP7 autoMARylation, leading to enhanced PARP7 stability and increased PARP7 protein levels, acts to drive adipogenesis.

### PARP7 regulates the proadipogenic gene expression program

Our results pointed to a role for PARP7 in promoting proadipogenic gene expression. To explore this possibility in more detail, we performed RNA sequencing in undifferentiated and differentiated (2 days) 3T3-L1 cells subjected to siRNA-mediated knockdown of *Parp7*. We confirmed the depletion of PARP7 protein by Western blotting (Suppl. Fig. S4A) and the knockdown of *Parp7* mRNA in the RNA-seq data (Suppl. Fig. S4B, *left*). We also observed the expected increase in *Cebpb* mRNA in the control sample during differentiation in the RNA-seq data (Suppl. Fig. S4B, *right*). Depletion of PARP7 had a profound effect on proadipogenic gene expression, abrogating the differentiation-induced up- or downregulation of over 2,000 genes (Fig. 4A; Suppl. Fig. S4C), including key adipogenic marker genes, such as *Adipoq* (Suppl. Fig. S4D). Gene ontology analyses of the affected genes revealed an enrichment of terms related to metabolism, especially lipid metabolism, and adipocyte differentiation (Fig. 4B, downregulated upon PARP7 depletion) and immune response genes (Suppl. Fig. S4E, upregulated upon PARP7 depletion). These results link the nuclear localization and differentiation-induced stability of PARP7 to proadipogenic gene expression outcomes.

**Figure 4.**
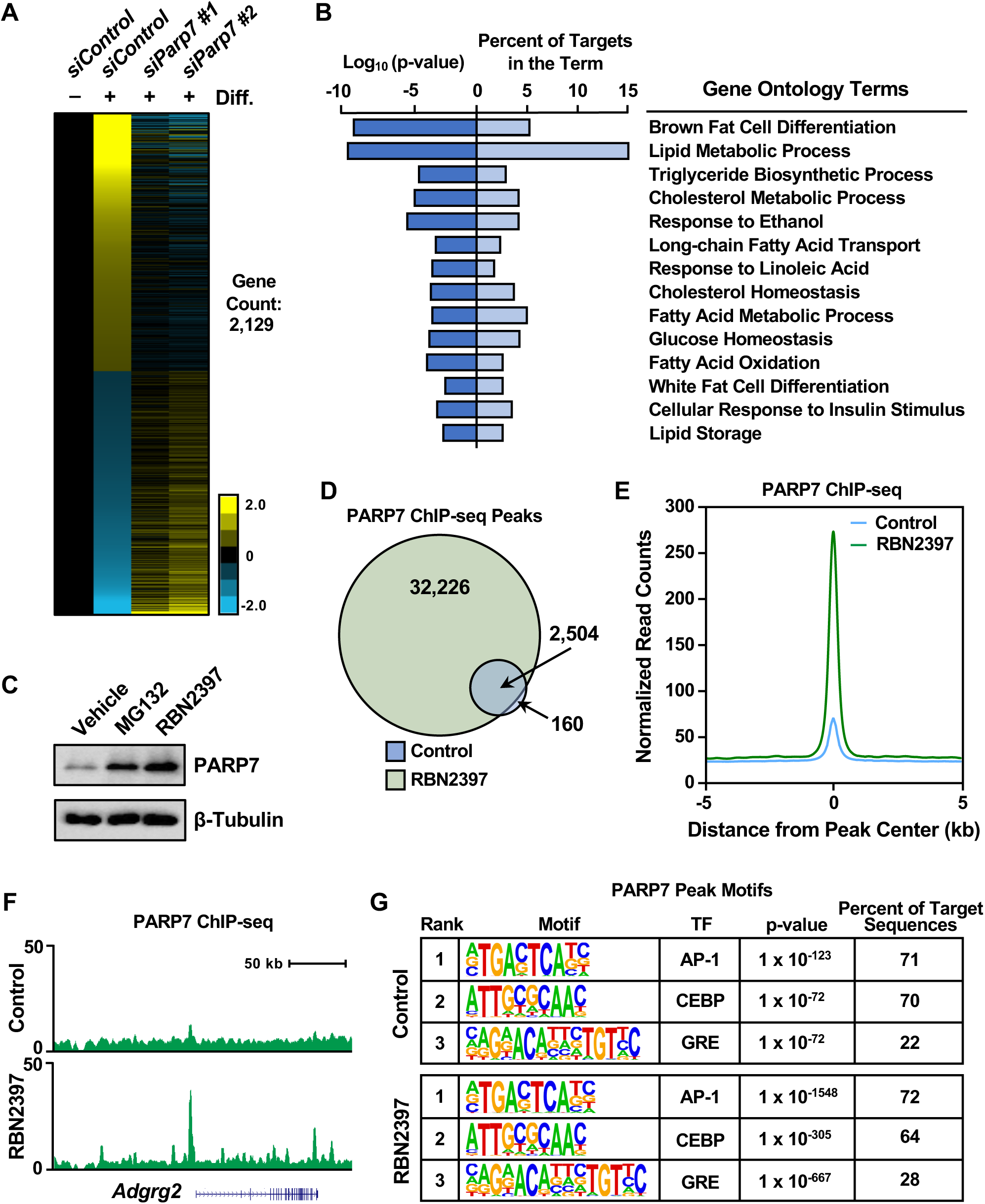
PARP7 regulates the proadipogenic gene expression program. (A) Heatmap of RNA-seq data from 3T3-L1 cells showing the effect of siRNA-mediated knockdown of *Parp7* at 2 days of differentiation using MDI cocktail. siRNA-mediated knockdown was performed two days before differentiation was initiated. (B) Gene Ontology terms for genes downregulated by *Parp7* knockdown after 2 days of differentiation. (C) Western blot showing PARP7 levels in 3T3-L1 cells treated with DMSO vehicle, 10 µM MG132, or 400 nM RBN2397 after at 8 hours of differentiation. β-tubulin was used as a loading control. (D) Venn diagram showing the overlap of significant PARP7 ChIP-seq peaks from vehicle- or RBN2397 (400 nM)-treated 3T3-L1 cells. (E) Metaplot of PARP7 ChIP-seq data from 3T3-L1 cells showing increased enrichment of PARP7 binding upon treatment with 400 nM RBN2397. The data are centered at the peak summits. (F) Genome browser tracks of PARP7 ChIP-seq data at the *Adgrg2* gene showing increased enrichment of PARP7 upon treatment with 400 nM RBN2397. (G) Representative motif enrichment at significant PARP7 ChIP-seq peaks in 3T3-L1 cells for the control *(top)* and 400 nM RBN2397 *(top)* treatment groups.

### PARP7 binds to chromatin and is required for the binding of C/EBPβ

Previous studies have shown that PARP7 can act as a cofactor for various transcription factors (Bindesboll et al., 2016a; Kamata et al., 2021b; MacPherson et al., 2013; Rasmussen et al., 2021). Knowing this, we asked if PARP7 is localized to chromatin during adipogenesis. To explore the possibility that PARP7 might bind to chromatin and serve as cofactor for the proadipogenic transcription factor C/EBPβ, we used chromatin immunoprecipitation (ChIP)- qPCR assays in 3T3-L1 cells. We observed that both PARP7 and C/EBPβ colocalized at the promoters of C/EBPβ target genes (e.g., *Pparg* and *Cebpb*), with the enrichment for both reduced considerably upon CRISPR/Cas9-mediated knockout of *Parp7* (Suppl. Fig. S5A).

To expand these results to a global scale, we performed ChIP-seq for PARP7 (at 8 hours of differentiation) and Cleavage Under Targets and Release Using Nuclease (CUT&RUN) (Skene and Henikoff, 2017) for C/EBPβ (at 24 hours of differentiation) in 3T3-L1 cells under various conditions, including siRNA-mediated knockdown of *Parp7* and treatment with RBN2397. The timing of the genomic localization assays was based on the timing of the expression maxima and functions of PARP7 and C/EBPβ. Because the half-life of PARP7 is very short (Fig. 2C) and treatment with RBN2397 stabilizes PARP7 (Fig. 4C), treatment with RBN2397 provided a unique opportunity to enhance the signal for PARP7, a notoriously difficult protein to analyze by ChIP-seq. Indeed, we observed stronger PARP7 binding and a 13-fold increase in the number of significant PARP7 peaks compared to untreated 3T3-L1 cells (Fig. 4, D through F). Importantly, the peaks observed in the untreated samples were almost completely a subset of the peaks observed in the in the RBN2397-treated cells (Fig. 4, D and F), indicating that the RBN2397 stabilizes bona fide PARP7 binding sites.

Many of the sites of PARP7 binding overlapped C/EBPβ binding, for example at key C/EBPβ target genes (e.g., *Pparg* and *Cebpa*) (Suppl. Fig. S5, B and C). PARP7 binding occurred primarily at enhancers and intergenic regions, but also at promoters and the first exon (Suppl. Fig. S5, D and E). Importantly, the genomic locations of PARP7 binding were similar ± RBN2397 treatment (Suppl. Fig. S5E). Moreover, the distance of the PARP7 peaks from the transcription start sites (TSSs) of PARP7-regulated genes was similar ± RBN2397 treatment (Suppl. Fig. S5, F and G). However, the distance of the PARP7 peaks from the TSSs of PARP7- regulated genes with MG132 treatment was generally farther than the distances observed ± RBN2397 treatment (Suppl. Fig. S5H). Finally, the sequences enriched under the PARP7 peaks ± RBN2397 treatment were similar, revealing motifs for C/EBPβ, AP-1 and related transcription factors, and glucocorticoid receptor (Fig. 4G; Suppl. Fig. S5, I and J). Together, these results indicate that we were able to successfully perform genomic localization analyses for PARP7, which linked PARP7 to C/EBPβ across the genome.

To expand on our observations, we examined the following in 3T3-L1 cells upon PARP7 depletion: (1) the genomic localization of C/EBPβ and histone H3 lysine 27 acetylation (H3K27ac) using CUT&RUN and (2) chromatin accessibility using assay for transposase accessible chromatin using sequencing (ATAC-seq) (Buenrostro et al., 2013) (Fig. 5A). We confirmed the depletion of PARP7 upon siRNA-mediated knockdown of *Parp7*, as well as the expression and phosphorylation of C/EBPβ (Fig. 5B). The C/EBPβ peaks in our CUT&RUN data matched well with previously reported C/EBPβ peaks from ChIP-seq data (Siersbaek et al., 2014) (Suppl. Fig. S6, A and B). At the key C/EBPβ target gene *Pparg*, we observed a dramatic reduction in the binding of C/EBPβ and enrichment of H3K27ac, with limited effects on chromatin accessibility upon depletion of PARP7 (Fig. 5C). On a global scale, the depletion of PARP7 caused a dramatic reduction in C/EBPβ at ∼70% of the >55,000 significantly called peaks, with ∼30% unaffected (“maintained”) and very few gained peaks (Fig. 5, D and E). The depleted C/EBPβ peaks localized predominantly at promoters, whereas the maintained peaks localized predominantly at enhancers (Suppl. Fig. S6C). The global results for H3K27ac and chromatin accessibility at the C/EBPβ peaks largely mirrored the results observed at the *Pparg* gene, with a dramatic reduction in the enrichment of H3K27ac and more modest reductions in chromatin accessibility (Suppl. Fig. S6, D through H).

**Figure 5.**
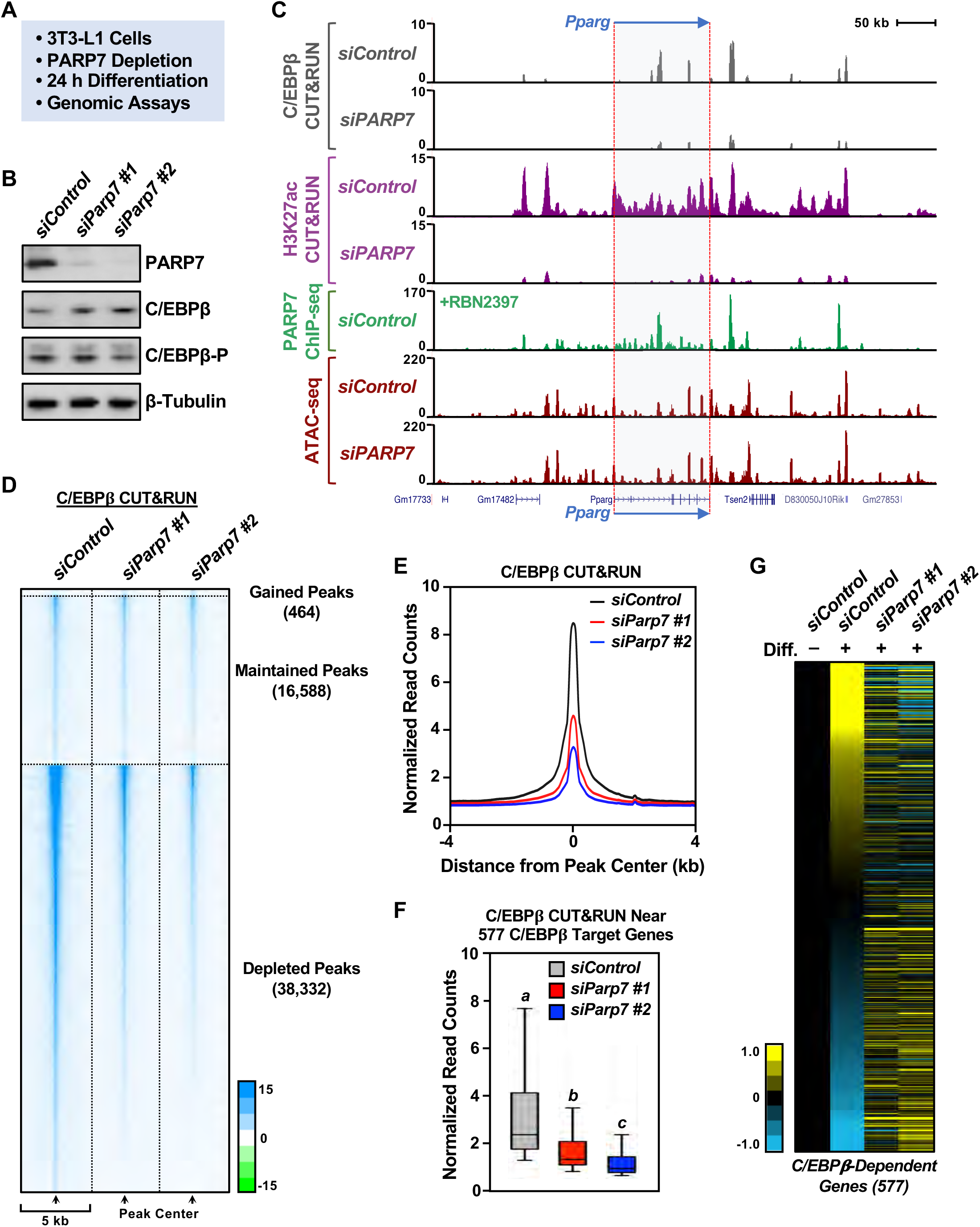
PARP7 binding to chromatin is required for the C/EBP**β**-dependent proadipogenic gene expression program. **(A)** Schematic of experimental set-up used for the genomic assays in 3T3-L1 cells. **(B)** Western blots showing the effects of *Parp7* knockdown on the levels of total C/EBPβ and phosphorylated C/EBPβ (C/EBPβ-P) at day 1 of differentiation in 3T3-L1 cells. The cells were subjected to siRNA-mediated *Parp7* knockdown two days before differentiation was induced using MDI cocktail. β-tubulin was used as a loading control. **(C)** PARP7 and C/EBPβ share genomic binding sites. Browser tracks of genomic data at the *Pparg* gene. 3T3-L1 cells were subjected to control or *Parp7* knockdown, followed by genomic assays: C/EBPβ and H3K27ac CUT&RUN, PARP7 ChIP-seq, and ATAC-seq. PARP7 ChIP- seq was performed after treatment with 400 nM RBN2397 to stabilize the PARP7 protein. **(D)** Heat map showing gained, maintained, and depleted C/EBPβ CUT&RUN peaks upon siRNA-mediated knockdown of *Parp7* in 3T3-L1 cells. **(E)** Metaplot of C/EBPβ CUT&RUN data centered at significant C/EBPβ peaks showing decreased C/EBPβ binding upon siRNA-mediated knockdown of *Parp7* in 3T3-L1 cells. **(F)** Box plot quantification of reads from C/EBPβ CUT&RUN peaks near 577 C/EBPβ target genes defined previously (Siersbaek et al., 2011) after siRNA-mediated knockdown of *Parp7* in 3T3-L1 cells. Bars marked with different letters are significantly different from each other. Wilcox Rank Sum test, p < 4.652 x 10^-15^. **(G)** RNA-seq heatmap showing the regulation of 577 C/EBPβ target genes defined previously (Siersbaek et al., 2011) in 3T3-L1 cells with siRNA-mediated knockdown of *Parp7*.

When we focused our analyses on 577 C/EBPβ target genes defined previously (Siersbaek et al., 2011), we observed a substantial enrichment of PARP7 binding within 20 kb of the promoters of the genes that was not observed for a random set of 577 unregulated genes (Suppl. Fig. S6I). We also observed a dramatic reduction in C/EBPβ enrichment at the nearest neighboring binding sites (Fig. 5F) and a concomitant alteration in the expression of the corresponding genes (Fig. 5G). Collectively, our genomic analyses provide evidence of a strong functional link between PARP7 and C/EBPβ that drives a proadipogenic gene expression program. Reduced autoMARylation and stabilization of PARP7 upon differentiation-induced depletion of nuclear NAD^+^ play a key role in driving this process.

### PARP7 regulates adipogenesis in vivo

To examine the broader implications of PARP7-mediated regulation of adipogenesis in vivo, we employed a *Parp7* knockout mouse model generated using the ‘knockout first’ approach described previously (Skarnes et al., 2011). In this model, the *Parp7^tm1a^* allele acts as a knockdown, rather than a true knockout, but is sufficient for functional analyses in vivo. In initial experiments, we examined the effects of whole-body knockdown of *Parp7* in response to 8 weeks on a high fat diet (*Parp7^WT/WT^* versus *Parp7^tm1a/tm1a^*) (Fig. 6A). We confirmed the depletion of PARP7 in the *Parp7^tm1a/tm1a^* mice by Western blotting (Fig. 6B). As expected, we observed an increase in the size, weight, and fat content of the *Parp7^WT/WT^*mice on the high fat diet, with a significant reduction in all three parameters in the *Parp7^tm1a/tm1a^* mice (Fig. 6, C through E; Suppl. Fig. S7, A through C). In primary preadipocytes isolated from SVF, we observed differentiation-induced and RBN2397-induced increases in PARP7 levels, as well as depletion of PARP7 in cells isolated from *Parp7^tm1a/tm1a^*mice (Suppl. Fig. S7, D and E). The depletion of PARP7 in primary preadipocytes isolated from *Parp7^tm1a/tm1a^* mice exhibited a reduction in the expression of differentiation-associated marker genes (e.g., *Adipoq* and *Fabp4*) and the accumulation of lipids relative to *Parp7^WT/WT^* mice (Suppl. Fig. S7, F through I). Together, these results demonstrate that PARP7 is required for the differentiation of adipocytes in vivo.

**Figure 6.**
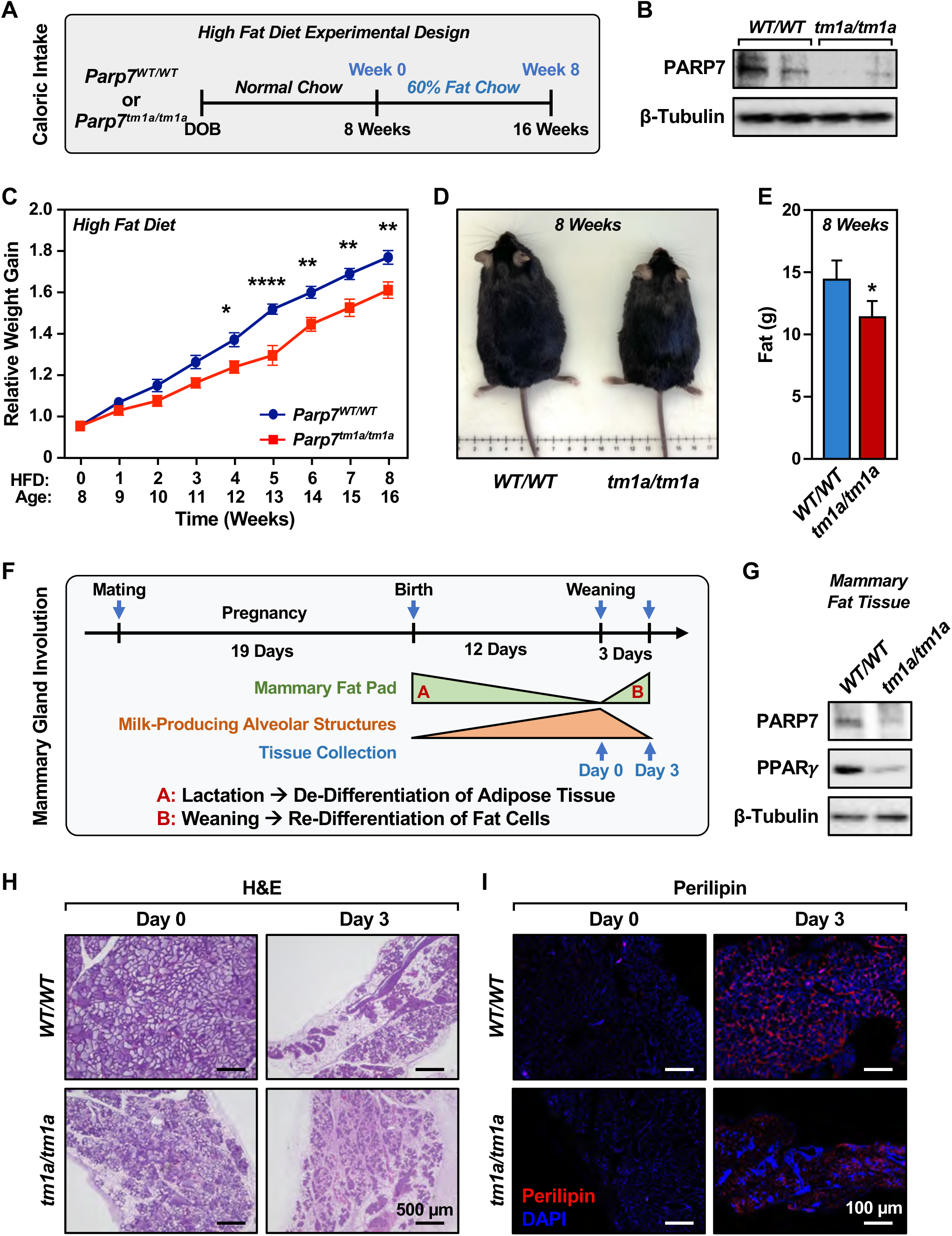
PARP7 is required for adipogenesis in vivo. **(A)** Schematic diagram showing the design of the high fat diet experiments in mice. **(B)** Western blot showing PARP7 levels in subcutaneous fat tissues isolated from *Parp7^WT/WT^* or *Parp7^tm1a/tm1a^* mice after 8 weeks on a high (60%) fat diet. β-tubulin was used as a loading control. **(C)** Line graph showing the relative weight gain of *Parp7^WT/WT^* or *Parp7^tm1a/tm1a^*mice over 8 weeks on a high fat diet. Each mouse was normalized to its own weight at the start of the high fat diet. Each point represents the mean ± SEM; n = 12. ANOVA; * = p < 0.0332, ** = p < 0.0021 , **** = p < 0.0001. **(D)** Representative images of *Parp7^WT/WT^* and *Parp7^tm1a/tm1a^* mice after 8 weeks of high fat diet. **(E)** Bar graph quantification of the fat content in *Parp7^WT/WT^* or *Parp7^tm1a/tm1a^*mice, as determined by MRI, after 8 weeks on a high fat diet. Each bar represents the mean + SEM; n = 12. The bar marked with an asterisk is significantly different from the control; Student’s t-test; * = p < 0.0332. **(F)** Schematic diagram showing the experimental design for the mouse mammary gland involution experiments. Adipogenesis was examined in the involuting mammary gland at day 0 and day 3 of involution in *Parp7^WT/WT^*or *Parp7^tm1a/tm1a^* mothers after the weaning of the pups (*Parp7^WT/tm1a^*) at day 12 post-partum. **(G)** Western blots showing PARP7 and PPARy levels in *Parp7^WT/WT^* or *Parp7^tm1a/tm1a^* mouse mammary fat tissues at day 3 of involution. β-tubulin was used as a loading control. **(H and I)** Structural changes and reduced fat storage in *Parp7^tm1a/tm1a^*mice versus *Parp7^WT/WT^* mice. Representative (H) H&E staining and (I) perilipin immunofluorescent staining of mammary fat pad tissues from *Parp7^WT/WT^* or *Parp7^tm1a/tm1a^* mice at day 0 and 3 of involution. DNA was stained with DAPI. Scale bars: (H) 500 µm and (I) 100 µm.

To explore the role of PARP7 in adipocytes populating a specific tissue, we examined mammary gland involution in *Parp7^WT/WT^* versus *Parp7^tm1a/tm1a^* mice. Previous studies have shown that the adipocyte population in the mammary fat pad in mice dedifferentiates as milk-producing alveoli form. After the pups are weaned, adipogenesis occurs to repopulate the mammary fat pad with adipocytes (Wang and Scherer, 2019) (Fig. 6F). We took advantage of this naturally-occurring process to determine if the depletion of PARP7 could impact repopulation of the mammary fat pad with adipocytes through the process of adipogenesis. As expected, both PARP7 and PPARγ levels were depleted in the *Parp7^tm1a/tm1a^* mice (Fig. 6G). The *Parp7^tm1a/tm1a^* mice showed gross morphological differences in the mammary gland compared to *Parp7^WT/WT^*mice (Fig. 6H; Suppl. Fig. S7, J and K). At 3 days post-weaning, increased staining for perilipin, a protein located on the surface of intracellular lipid droplets, indicated the repopulation of the mammary fat pad with adipocytes in *Parp7^WT/WT^* mice during involution, an effect that was not observed in *Parp7^tm1a/tm1a^* mice (Fig. 6I; Suppl. Fig. S7L). These results provide additional evidence indicating that PARP7 is required for the differentiation of adipocytes in vivo.

### PARP7 is required for maintaining the nutrient content of mother’s milk and the viability of pups

During the course of breeding the *Parp7* knockout, we observed an increased length of gestation (by ∼2 days) in the *Parp7^tm1a/tm1a^*mice versus *Parp7^WT/WT^* mice (Fig. 7A). In addition, we observed a dramatic decrease in the number of viable pups at 1 day post-partum (Fig. 7B), which skewed the breeding ratios from *Parp7^WT/tm1a^*x *Parp7^WT/tm1a^* crosses (Suppl. Fig. S8A). Given the dramatic decrease in the levels of PPARγ expression that we observed in mammary adipose tissues from female *Parp7^tm1a/tm1a^*mice (Fig. 6G), together with previous literature linking PPARγ to milk lipid regulation and the quality of mother’s milk (Wan et al., 2007; Yang et al., 2018), we considered the possibility that the lethality of the pups might be related to altered nutrient content in the milk from the *Parp7^tm1a/tm1a^* mice. To address this, we performed metabolomics for lipids and polar metabolites in the mammary tissue fat pads and milk from *Parp7^WT/WT^*and *Parp7^tm1a/tm1a^* mice. Although some lipids showed increased levels in the mammary fat pad from *Parp7^tm1a/tm1a^*mice, most lipids that exhibited a significant change versus *Parp7^WT/WT^* were decreased (Fig. 7C), with the distribution of lipid types indicated in Fig. 7D. In contrast, the levels of the individual lipid species identified in milk from *Parp7^tm1a/tm1a^* mice were variable. Although we did not observe statistically significant reduction of any individual lipid (Suppl. Fig. S8B), the total fatty acid content in milk was significantly reduced as indicated by the top 50 lipids that we assayed (Fig. 7E). We did not observe significant changes in 15(S)- hydroxyeicosatetraenoic acid, [15(S)-HETE, an active metabolite of arachidonic acid] or 13- hydroxyoctadecadienoic acid (13-HODE, a fatty acid metabolite produced from linoleic acid) (Suppl. Fig. S8C), which have been implicated previously in inflammatory “toxic” milk from *Pparg* knockout mice (Wan et al., 2007).

**Figure 7.**
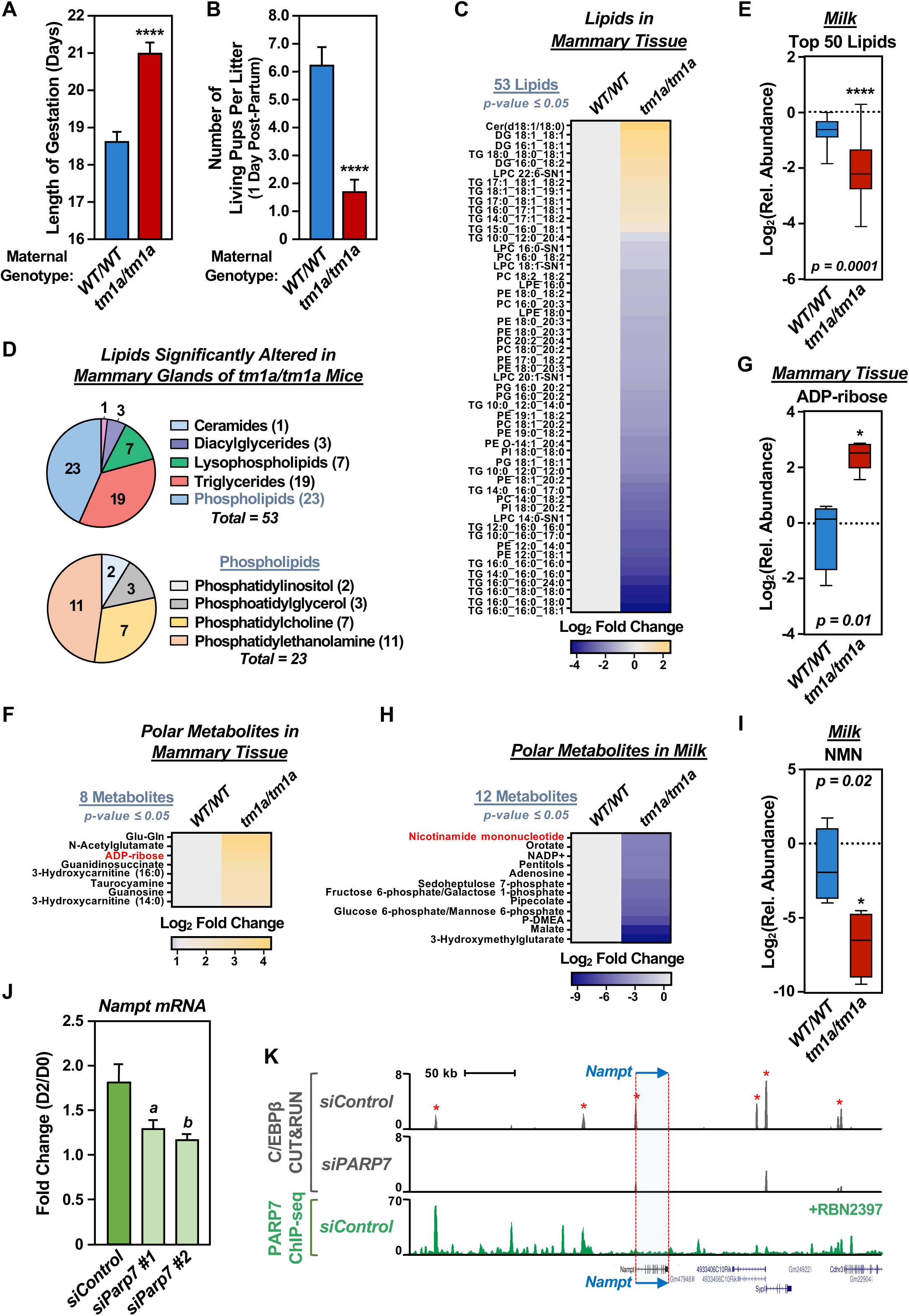
Altered adipose tissue metabolism and milk nutrient content in *Parp7^tm1a/tm1a^* mice. **(A and B)** Altered length of gestation and pup survival in *Parp7^tm1a/tm1a^*versus *Parp7^WT/WT^* female mice. *Parp7^WT/WT^* and *Parp7^tm1a/tm1a^* female mice were subjected to timed matings. The length of gestation (A) and the number of pups surviving at one day post-partum (B) were assessed. Each bar represents the mean + SEM; n = 7. Unpaired t-test; **** = p < 0.0001. **(C through E)** Lipidomics analysis in mammary fat pads and milk from *Parp7^WT/WT^* and *Parp7^tm1a/tm1a^* female mice (mammary fat pads, n = 4 mice for WT, n = 5 mice for tm1a; milk, n = 4 mice for each group). Samples were collected at the time of weaning (day 12 post-partum, day 0 of involution). The samples were subjected to mass spectrometry-based lipidomics. (C) Heat map showing changes in 53 lipids in mammary fat pads that were significantly altered (p ≤ 0.05) in *Parp7^tm1a/tm1a^* versus *Parp7^WT/WT^* mice. (D) Pie charts showing changes in different lipid types significantly altered in the mammary glands of *Parp7^tm1a/tm1a^* mice from panel (C). (E) Box plot showing the average change in the top 50 altered lipids in milk in aggregate. Unpaired t-test; **** p < 0.0001. **(F through I)** Metabolomics analysis in mammary fat pads and milk from *Parp7^WT/WT^*and *Parp7^tm1a/tm1a^* mice (mammary fat pads, n = 4 mice for WT, n = 5 mice for tm1a; milk, n = 4 mice for each group). Samples were collected at the time of weaning (day 12 post-partum, day 0 of involution). The samples were subjected to mass spectrometry-based metabolomics. (F and H) Heat maps showing changes in metabolites in (F) mammary fat pads and (H) milk that were significantly altered (p ≤ 0.05) in *Parp7^tm1a/tm1a^*versus *Parp7^WT/WT^* mice (8 out of 8 shown for mammary tissue; 12 out of 64 shown for milk). (G and I) Box plots showing changes in (G) ADP-ribose levels in mammary fat pads and (I) nicotinamide mononucleotide (NMN) levels in milk from *Parp7^tm1a/tm1a^* versus *Parp7^WT/WT^* mice. Unpaired t-test with Welch’s correction;* p = 0.01 or 0.02, as indicated. **(J and K)** Regulation of *Nampt* gene expression by PARP7 in 3T3-L1 cells. (J) Effect of PARP7 depletion on *Nampt* gene expression determined by RNA-seq. 3T3-L1 cells were subjected to knockdown with control or *Parp7* siRNAs, followed by differentiation with MDI cocktail for 0 or 2 days. RNA-seq data for the *Nampt* gene were collected, normalized to *Gapdh*, and expressed as fold change (day 2 versus day 0). Each bar represents the mean + SEM; n = 3. Bars with different letters are significantly different; Unpaired t-test (two tailed), p-values for *a* and *b* are p < 0.0455 and p < 0.0179, respectively. (K) Browser tracks of genomic data at the *Nampt* gene. 3T3-L1 cells were subjected to control or *Parp7* knockdown, followed by genomic assays: C/EBPβ CUT&RUN and PARP7 ChIP-seq (+ 400 nM RBN2397 to stabilize the PARP7 protein). Asterisks indicate C/EBPβ peaks that are reduced upon PARP7 depletion.

We also examined polar metabolites in the mammary tissue fat pads and milk from *Parp7^WT/WT^* and *Parp7^tm1a/tm1a^* mice. We observed a significant increase in 8 polar metabolites from the mammary fat pads of *Parp7^tm1a/tm1a^* mice versus *Parp7^WT/WT^* mice (Fig. 7F; Suppl. Fig. S8D), including ADP-ribose (Fig. 7G). In contrast, we observed a significant decrease in 64 polar metabolites from the milk of *Parp7^tm1a/tm1a^*mice versus *Parp7^WT/WT^* mice (Fig. 7H; Suppl. Fig. S8E), including NMN (Fig. 7I). The decrease in NMN was supported by the results of pathway analyses (Suppl. Fig. S8, F and G). Given the dramatic reduction of NMN in the milk from *Parp7^tm1a/tm1a^* mice, we considered the possibility that PARP7 plays a direct role in the regulation of *Nampt*, the gene encoding nicotinamide phosphoribosyltransferase (NAMPT) – the enzyme that synthesizes NMN. Indeed, we observed that the differentiation-induced increase in *Nampt* expression observed in 3T3-L1 cells is impaired with PARP7 depletion (Fig. 7J). Moreover, the *Nampt* gene has a number of PARP7 binding sites, as well as a collection of PARP7-dependent C/EBPβ binding sites, sites located nearby (Fig. 7K).

Collectively, these results demonstrate that lipid production is impaired in the mammary fat pads of *Parp7^tm1a/tm1a^* mice, which likely accounts for the reduced lipid content in the milk of *Parp7^tm1a/tm1a^* mice. Moreover, the levels of NMN, a key nutrient in mother’s milk (Hattori et al., 2024; Saito et al., 2023), is dramatically reduced in the milk of *Parp7^tm1a/tm1a^*mice, which may contribute to the poor survival of the pups. Finally, gene expression and genomic analyses link the PARP7-dependent C/EBPβ-mediated proadipogenic gene expression program directly to the regulation of NMN production via the expression of *Nampt*.

## Discussion

Adipogenesis is a complex process in which cellular signals drive a carefully orchestrated series of molecular events that lead to proadipogenic gene expression and, ultimately, the intracellular accumulation of lipids. In the studies described herein, we demonstrate that PARP7, a MART in the PARP family, serves as a key nuclear regulatory protein during adipogenesis. Interestingly, the functions of PARP7 in this system are independent of its ability to transmodify substrate proteins. Rather, PARP7 serves as a coregulator of the proadipogenic transcription factor, C/EBPβ, after PARP7 is auto-stabilized in response to decreasing nuclear NAD^+^ levels. We introduce ‘NAD^+^ sensing’ to explain and integrate the series of events that lead from proadipogenic signaling and reduced nuclear NAD^+^ levels to PARP7 stabilization and coactivation of C/EBPβ. These studies enhance our understanding of the molecular mechanisms that control adipogenesis.

### PARP7 is required for adipogenesis

Our siRNA screen of nuclear and cytosolic MARTs in 3T3-L1 preadipocytes using a variety of endpoint assays identified PARP7 as a robust regulator required for efficient adipogenesis. These results were extended to a mouse genetic model with whole body depletion of PARP7, where we observed (1) reduced weight gain and fat accumulation on a high fat diet and (2) a failure to repopulate the lactating mammary gland with adipocytes during post-weaning involution. Thus, in three different biological models of adipogenesis and for multiple different molecular, cellular, and tissue endpoint assays (i.e., fat accumulation, gene expression, metabolite levels), we observed PARP7 to be required for adipogenesis. Taken together with our previous studies identifying PARP1 as a molecular regulator of adipogenesis (Huang et al., 2020; Luo et al., 2017; Ryu et al., 2018), these results highlight the growing role of PARP family members in the control of adipogenesis, albeit through distinct and complementary mechanisms. PARylation by PARP1 inhibits the DNA binding and transcriptional activities of C/EBPβ (Luo et al., 2017), whereas autoMARylation promotes the degradation of PARP7, preventing it from serving as a coactivator of C/EBPβ. As described below, these molecular events are driven by NAD^+^ sensing mechanism that underlie the biochemical and molecular events leading to adipogenesis.

### PARP7 serves as a transcriptional coregulator of C/EBPβ

Multiple lines of investigation from our study support the conclusion that PARP7 serves as a coregulator of C/EBPβ. Both PARP7 and C/EBPβ drive proadipogenic gene expression, with significantly overlapping target gene sets. Moreover, both PARP7 and C/EBPβ share significantly overlapping binding sites across the genome, and PARP7 binding sites are enriched for C/EBPβ motifs in the genomic DNA. Finally and most importantly, depletion of PARP7 causes a dramatic reduction in the enrichment of C/EBPβ at about 70 percent of its binding sites. While we have not yet worked out the specific mechanistic details for PARP7-mediated transcriptional coregulation of C/EBPβ, we note that depletion of PARP7 also causes a dramatic reduction in the levels of H3K27ac, a mark associated with active chromatin, without dramatically impacting chromatin accessibility. p300, the major H3K27 acetyltransferase, has been shown previously to be required for the transcriptional regulation of C/EBPβ (Guo et al., 2015; Mink et al., 1997; Schwartz et al., 2003; Steger et al., 2010). A recent study has shown that PARP7 interacts with the transcription factor IRF3 and inhibits its interaction with CBP/p300 (Jeltema et al., 2025). Future studies will examine potential functional connections between chromatin binding by PARP7 and C/EBPβ, H3K27ac, and p300.

### PARP7 senses nuclear NAD^+^ concentrations to control gene expression

We have previously demonstrated and quantified a rapid and dramatic reduction in nuclear NAD^+^ concentrations within the first 4-8 hours after the signal-induced differentiation of preadipocytes (Ryu et al., 2018). They range from a high of ∼100 µM in undifferentiated preadipocytes to a low of ∼40 µM after 8 hours of differentiation. The Km of PARP7 for NAD^+^ is ∼70 µM, which should support the catalytic activity of PARP7 at the high nuclear NAD^+^ concentrations available pre-differentiation and reduce its catalytic activities at the low nuclear NAD^+^ concentrations available post-differentiation. This is what we observed experimentally. We have previously reported similar results for PARP1 (Ryu et al., 2018), which has a Km for NAD^+^ of ∼85 µM (Langelier et al., 2010). Thus, at the relatively high nuclear NAD^+^ concentrations in undifferentiated preadipocytes, PARP7 and PARP1 are active and can ADPRylate substrates that allow the control of adipogenesis – automodification in the case of PARP7 and transmodification of C/EBPβ and H2B in the case of PARP1 (Huang et al., 2020; Luo et al., 2017; Ryu et al., 2018). At the relatively low nuclear NAD^+^ concentrations 8 hours post-differentiation, these ADPRylation events decrease dramatically.

NAD^+^ sensing by PARP7 and PARP1 creates an NAD^+^-dependent switch from PARP1 inhibition of C/EBPβ to PARP7 coactivation of C/EBPβ, allowing the proadipogenic gene expression program to proceed. Previous studies demonstrated that the transcriptional corepressor CtBP can function as a nuclear redox sensor during redox-regulated transcription by virtue of a much greater affinity for NADH than NAD^+^ (Fjeld et al., 2003; Zhang et al., 2006). NAD^+^ sensing by PARP7 and PARP1 is similar in some respects, but differs in that NAD^+^ is a substrate that is consumed by PARP7 and PARP1, which may contribute to the reduction in NAD^+^ levels as differentiation proceeds. Moreover, NAD^+^ sensing by these PARPs is likely to occur independently of the redox state in the cell.

### AutoMARylation regulates PARP7 stability through an E3 ligase-ubiquitin-proteasome pathway

Our results demonstrate that in undifferentiated preadipocytes, PARP7 is highly unstable and has a half-life of minutes, similar to what has been reported in prostate cancer cells (Kamata et al., 2021a). This instability is mediated by a mechanism requiring autoMARylation, ubiquitylation, and proteasome-mediated degradation of PARP7 when nuclear NAD^+^ concentrations are sufficiently high. We identified DTX2 and RNF114 as E3 ligases that promote the ubiquitin-mediated degradation of PARP7. Interestingly, both DTX2 and RNF114 contain “reader” domains known to interact with ADP-ribose (Li et al., 2023; Munzker et al., 2024). A series of recent studies have shown that Deltex E3 ligases can ubiquitylate protein-linked ADPR, specifically at the 3’ hydroxyl of the adenosine moiety, generating a noncanonical ubiquitin ester-linked species (Bejan et al., 2025; Chatrin et al., 2020; Kelly et al., 2024; Zhu et al., 2022). Given these observations, it remains formally possible that PARP7 is ubiquitylated at the sites of autoMARylation. They also raise the interesting possibility that different combinations of E3 ligases with distinct ADPR and ADPR-ubiquitin reading activities facilitate the ubiquitylation and proteasome-mediated degradation of autoMARylated PARP7.

### PARP7 controls the metabolome in lactating mammary glands and milk

Our genetic and metabolomic analyses of the mammary gland fat pads and milk from lactating female mice have revealed critical biological roles for PARP7. The dramatic reduction in a wide array of lipids in the mammary gland fat pads and milk upon genetic depletion of PARP7 highlight the critical role of PARP7 in adipogenesis in these organs. In addition to the changes in lipids, we also observed a significant increase in ADPR levels in mammary gland fat pads and a significant decrease in NMN in the milk from the PARP7 depleted mice. The increase in ADPR levels in lactating mammary glands upon PARP7 depletion may be due to turnover of nuclear PAR that accumulates in undifferentiated preadipocytes, prior to differentiation-induced reductions in nuclear NAD^+^. The decrease in ADPR levels in the milk from lactating mammary glands is likely due, at least in part, to reduced expression of *Nampt*, the gene encoding NAMPT, the enzyme that produces NMN, which is a direct transcriptional target of PARP7 and C/EBPβ. NMN is a key nutrient in mother’s milk (Hattori et al., 2024; Saito et al., 2023), and its reduced levels in the milk of *Parp7^tm1a/tm1a^* mice may contribute to the poor survival of the pups.

Collectively, our results extend the biology of PARP7 beyond its previously recognized function as a stress-responsive regulatory protein in pluripotency, neuronal function, immune responses and immunity, and oncology to adipogenesis and perinatal health. Moreover, our results describe the molecular events that regulate these downstream biological functions.

## Materials and Methods

### Antibodies, chemicals, and specialized reagents

The custom rabbit polyclonal antiserum against PARP7 was generated by Pocono Rabbit Farm and Laboratory by using a purified recombinant antigen comprising the amino-terminal half of PARP7 (amino acids 1-324). The serum was then purified using antigen affinity chromatography.

Other antibodies used were as follows: PARP7 (Thermo Fisher Scientific, PA5-40774; RRID:AB_2607074), C/EBPβ (Invitrogen, PA5-120052; RRID:AB_2913624, Invitrogen, PA5-86117; RRID:AB_2802916, and Cell Signaling, 3082; RRID:AB_2260365), Phospho-C/EBPβ (Thr235) (Cell Signaling, 3084S; RRID:AB_2260359), PPARγ (81B8) (Cell Signaling, 2443; RRID:AB_823598), DTX2 (ThermoFisher, PA5-109664; RRID:AB_2855075), RNF114 (Proteintech, 14338-1-AP; RRID:AB_3085435), Histone H3K27ac (Active Motif, 39134; RRID:AB_2722569), FLAG (Sigma-Aldrich, F3165; RRID:AB_259529), MAR binding reagent (Millipore, MABE1076; RRID:AB_2665469), PAR binding reagent (Millipore, MABE1031; RRID:AB_2665467), Perilipin (Biosynth, 20R-PP004; RRID: AB_3665667), Ubiquitin (E4I2J) (Cell Signaling, 43124; RRID:AB_2799235), K48-linkage specific polyubiquitin (Cell Signaling, 4289; RRID:AB_10557239), and β-tubulin (Abcam, ab6046; RRID:AB_2210370). Secondary antibodies included Goat anti-rabbit HRP-conjugated IgG (ThermoFisher, 31460; RRID:AB_228341), Goat anti-mouse HRP-conjugated IgG (ThermoFisher, 31430; RRID:AB_10960845), Rabbit IgG (ThermoFisher, 10500C; RRID:AB_2532981), and Alexa Fluor 594 donkey anti-rabbit IgG (ThermoFisher, A-21207; RRID:AB_141637).

For PARP7 inhibition, we used RBN2397 (synthesized by Acme Bioscience) (Gozgit et al., 2021). Other chemicals for cell treatments included: Doxycycline (Sigma, D9891), MG132 (Sigma, M7449), cycloheximide (Sigma, C7698), β-nicotinamide mononucleotide (NMN) (Sigma Cat, N350), insulin (Sigma, I5500), 3-isobutyl-1-methylxanthine (Sigma, 410957), and dexamethasone (Sigma, D4902).

### Cell culture and treatments

#### Isolation of primary preadipocytes from the stromal vascular fraction (SVF) of white adipose tissue

SVF cells were isolated as described previously (Gupta et al., 2012). Briefly, 4 to 6-week-old male mice were sacrificed and the inguinal white adipose tissue (WAT) was collected. The WAT was washed, pooled, minced, and digested for 2 hours at 37°C in 10 ml of digestion solution [100 mM HEPES pH 7.4, 120 mM NaCl, 50 mM KCl, 5 mM glucose, 1 mM CaCl_2_, 1 mg/mL collagenase D (Roche, 11088858001), and 1.5% BSA]. The digested WAT tissue was filtered through a 100 μm cell strainer to remove undigested tissue, and 30 ml of SVF cell culture medium [10% FBS, 1% penicillin/streptomycin in DMEM/F12, GlutaMAX (Life Technologies, 10565-018)] was added to dilute the digestion buffer. The flow-through was centrifuged for 5 minutes at 600 x *g* to collect the SVF cells. The cell pellet was resuspended in 10 ml of SVF culture medium, and passed through a 40 μm cell strainer to remove clumps of cells and large adipocytes. The cells were collected again by centrifugation at 600 x *g* for 5 minutes, resuspended in SVF culture medium (5 ml per 2 mouse equivalents), and plated in a 6 cm diameter collagen-coated culture dish until well attached.

#### Cell culture

VF cells (Rodeheffer et al., 2008; Van et al., 1976) were grown in SVF culture medium until confluent and were then cultured for 2 more days under contact inhibition. For differentiation, the cells were then treated for 2 days with an adipogenic cocktail (MDI), including 0.25 mM IBMX (3-isobutyl-1-methylxanthine; Sigma, 410957), 1 μM dexamethasone (Sigma, D4902), and 10 μg/mL insulin (Sigma, I5500). Subsequently, the cells were cultured in medium containing 10 μg/mL insulin for the indicated times before collection.

3T3-L1 cells (Green and Meuth, 1974) were obtained from the American Type Cell Culture (ATCC, CL-173; RRID:CVCL_0123). They were maintained in DMEM (Cellgro, 10- 017-CM) supplemented with 10% fetal bovine serum (Atlanta Biologicals, S11550) and 1% penicillin/streptomycin. For the induction of adipogenesis, the 3T3-L1 cells were grown to confluence and then cultured for 2 more days under contact inhibition. The cells were then treated for 2 days with an MDI adipogenic cocktail containing 0.25 mM IBMX, 1 μM dexamethasone, and 10 μg/mL insulin. Subsequently, the cells were cultured in medium containing 10 μg/mL insulin for the indicated times before collection.

HEK 293T cells were obtained from the ATCC (CRL-3216; RRID:CVCL_0063). They were maintained in DMEM (Cellgro, 10-017-CM) supplemented with 10% fetal bovine serum and 1% penicillin/streptomycin.

For all cell lines, fresh cell stocks were regularly replenished from the original stocks, authenticated for cell type identity using the GenePrint 24 system (Promega, B1870), and confirmed as *Mycoplasma*-free every three months using the Universal Mycoplasma Detection Kit (ATCC, 30-1012K).

#### Cell treatments

3T3-L1 cells were exposed to various treatments and culture conditions for the experiments described herein. For treatment with MG132 (Sigma, M7449), the cells were grown as described above, cells were then treated with 10 µM MG132 for 6 hours before collection. For treatment with RBN2397, upon addition of differentiation cocktail, 3T3-L1 cells were treated with 400 nM RBN2397. The treatment continued every other day with media changes until specified collection time. For PARP7 MAR immunoprecipitation assays, the cells were treated with RBN2397 for 16 hours. For treatment with NMN, the cells were grown as described above and were then treated with 10 mM of NMN upon addition of differentiation cocktail.

### Oil Red-O staining

3T3-L1 cells were cultured in 6- or 12-well plates and differentiated as described above. After 8 days of differentiation, the cells were rinsed twice with 1x phosphate-buffered saline (PBS) and fixed with 4% paraformaldehyde. The fixed cells were washed with water and incubated in 60% isopropanol for 5 minutes. After incubation, the isopropanol was removed and replaced with 0.3% Oil Red-O working solution for 5 minutes. The Oil Red-O working solution was prepared by diluting a stock solution (0.5% in isopropanol; Sigma, O1391) with water (3:2). Destaining and quantification was done by adding 500 µL of 100% isopropanol and incubating for 15 minutes on platform shaker, 200 µL was then transferred to a 96-well clear plate and analyzed using a plate reader at 550 nm.

### BODIPY staining

3T3-L1 cells were seeded into 4-well chambered slides (Thermo Fisher, 154534) and were grown and differentiated as described above. The cells were washed twice with PBS, fixed in 4% methanol-free paraformaldehyde for 15 minutes at room temperature, and washed three times with PBS. The fixed cells were stained with 1 μg/mL of BODIPY 493/503 (Life Technologies, D3922) for 10 minutes. The cells were washed three times with PBS and coverslips were placed on cells using the VectaShield Antifade Mounting Medium with DAPI (Vector Laboratories, H-1200) and images were acquired using a Keyence BZ-X810 fluorescence microscope.

### Immunofluorescent staining of cultured cells

3T3-L1 cells were seeded into 4-well chambered slides (Thermo Fisher, 154534) and were grown and differentiated as described above. The cells were washed twice with PBS, fixed in 4% methanol-free paraformaldehyde for 15 minutes at room temperature, and washed three times with PBS. The cells were permeabilized for 5 minutes using permeabilization buffer (PBS containing 0.1% Triton X-100), washed three times with PBS, and incubated for 1 hour at room temperature in Blocking Solution (PBS containing 1% BSA, 10% FBS, 0.3 M Glycine and 0.1% Tween-20). The fixed cells were incubated with a PARP7 antibody at a 1:1000 dilution in PBS at room temperature for 1 hour. The cells were washed three times with PBS and then incubated with Alexa Fluor 594 donkey anti-rabbit IgG (ThermoFisher, A-21207; RRID:AB_141637) at a 1:500 dilution in PBS for 1 hour at room temperature. The cells were washed three times with PBS and coverslips were placed on cells using the VectaShield Antifade Mounting Medium with DAPI (Vector Laboratories, H-1200). Images were acquired using a Keyence BZ-X810 fluorescence microscope.

### Cloning and plasmid construction for mouse PARP7 expression

***cDNA library.*** cDNA pools were prepared by extraction of total RNA from 3T3-L1 cells (mouse) using TRIzol Reagent (Invitrogen, 15596026), followed by reverse transcription using SuperScript III Reverse Transcriptase (Invitrogen, 18080093) with random hexamer primers (Roche, 11034731001) according to the manufacturer’s instructions.

***Generation and site-directed mutagenesis (SDM) of a Parp7 cDNA.*** The cDNA pools were used to amplify mouse *Parp7* cDNA for subsequent cloning using primers listed below. cDNA encoding N-terminally FLAG epitope-tagged mouse *Parp7* was cloned into *BamHI-* and *NotI*-digested pcDNA3 (Invitrogen, V79020; RRID:Addgene_128034) using the primers listed below. Catalytic-dead site point mutant for *Parp7* was generated by site-directed mutagenesis in the pcDNA3-FLAG-PARP7 vector using Pfu Turbo DNA polymerase (Agilent, 600250) with primers listed below which mutated the tyrosine at 564 to an alanine (pcDNA3-FLAG-PARP7- Y564A).

***Plasmid vectors for expression in bacteria.*** For making the N-terminal antigen for antibody production the sequence corresponding to the first 324 N-terminal amino acids of mouse PARP7 were amplified by PCR from pcDNA3-FLAG-PARP7 (described above). The product was then digested with *BamHI*- and *XhoI*- and were ligated into *BamHI*- and *XhoI*- digested pET19b (Novagen, 69677).

***Bacmid vectors for expression in insect cells.*** FLAG-tagged mouse PARP7 cDNA was amplified from pCDNA3-FLAG-PARP7 as described above. The PCR product was digested with *BamHI and EcoRI*, then ligated into a *BamHI*- and *EcoRI*-digested pFastBac1 plasmid. The recombinant pFastBac1 bacmid was then prepared for transfection into Sf9 cells by transformation into the DH10BAC *E. coli* strain with subsequent blue/white colony screening using the Bac-to-Bac system (Invitrogen) according to the manufacturer’s instructions.

***Lentiviral vectors for expression in mammalian cells*.** To generate Dox-inducible lentiviral vectors for expression of mouse wild-type or mutant mouse PARP7, the respective wild-type and Y564A mutant FLAG epitope-tagged cDNAs were amplified from the pcDNA3 expression vectors (described above). The cDNAs were cloned into *NheI*- and *XhoI*-digested pInducer20 (Addgene, 44012; RRID:Addgene_44012) using a Gibson Assembly kit (NEB, E2621) using the primers listed below.

***Primers used for cloning.*** The following cloning primer sequences were used:

- Amplification of *Parp7* from cDNA:

- *musParp7* Forward: 5’-ATGGAAGTGGAAACCACTGAACCTGAGC-3’

- *musParp7* Reverse: 5’-GTCAGTAACACTGTTTCCATTTAA-3’

- Cloning *Parp7* into pcDNA3:

- Forward: 5’-CTGGCTAGCGTTTAAACTTAATGGATTATAAGGATGACG-3’

- Reverse: 5’-AGCGGGTTTAAACGGGCCCTTTAAATGGAAACAGTGTTACTG-3’

- SDM Primers for *Parp7* in pcDNA3:

- *Parp7* Y564A Forward:

5’-AGCTTGCCTTCTTTGCGAAAGCACTGCCTTGTCCAAACATTG-3’

- *Parp7* Y564A Reverse:

5’-CAATGTTTGGACAAGGCAGTGCTTTCGCAAAGAAGGCAAGCT-3’

- Cloning *Parp7* N-terminus into pET19b:

- *Parp7-N* Forward:

5’-CCATGGCTCGAGCGCATCACCATCATCACCATGAAGTGGAAACCACTG AACCT-3’

- *Parp7-N* Reverse: 5’-ATGACTGTATTTGAAACAACTTGATAGGGATCCCGGG-3’

- Cloning *Parp7* into pInducer20:

- Forward: 5’-TCCGCGGCCCCGAACTAGTGATGGATTATAAGGATGACG-3’

- Reverse: 5’-GTTTAATTAATCATTACTACTTAAATGGAAACAGTGTTACTG-3’

### Generation of CRISPR/Cas9 lentiviral vectors for sgRNA expression and genome editing

The CRISPR plasmid lentiCRISPR v2 was obtained from Addgene (52961; RRID:Addgene_52961). We used the Dharmacon CRISPR Design Tool to design optimized single guide RNA (sgRNA) sequences for *Parp7*. A non-targeting sequence from the GeCKOv2 Mouse Library Pool A was used as the control (Sanjana et al., 2014). To clone the guide target sequence into lentiCRISPR v2, oligo pairs for each *Parp7* guide were synthesized. Each pair of oligos was annealed, diluted, and then assembled into lentiCRISPR v2 using Golden-Gate sgRNA cloning protocol (Sanjana et al., 2014; Shalem et al., 2014).

***sgRNA sequences.*** The following sgRNA target sequences were used:

- Parp7-1 (target in exon 2): AGTTAATCACATCATGGAAG

- Parp7-2 (target in exon 3): CTGAATTTGACCAACTACGA

- Parp7-3 (target in exon 5): TGCATATAAACCCACGTTTC

- Non-targeting: GCGAGGTATTCGGCTCCGCG

***Primers used for cloning.*** The following oligo pairs were used for cloning:

- Parp7 1 Forward: 5’-CACCGAGTTAATCACATCATGGAAG-3’

- Parp7 1 Reverse: 5’-AAACCTTCCATGATGTGATTAACTC-3’

- Parp7 2 Forward: 5’-CACCGCTGAATTTGACCAACTACGA-3’

- Parp7 2 Reverse: 5’-AAACTCGTAGTTGGTCAAATTCAGC-3’

- Parp7 3 Forward: 5’-CACCGTGCATATAAACCCACGTTC-3’

- Parp7 3 Reverse: 5’-AAACGAAACGTGGGTTTATATGCAC-3’

- Non-targeting oligo 1: 5’-CACCGGCGAGGTATTCGGCTCCGCG-3’

- Non-targeting oligo 2: 5’-AAACCGCGGAGCCGAATACCTCGCC-3’

### Expression and purification of recombinant PARP7 N-terminus, and generation of a polyclonal antiserum

Bacteria induced using IPTG was pelleted and lysed with lysis buffer (50 mM Na_2_HPO_4_ pH 4.0, 0.3 M NaCl, 1 mM PMSF, 8 M Urea). Lysate was sonicated, centrifuged to clear debris, and supernatant was incubated with Ni-NTA agarose beads. After incubation beads were washed 3 times with wash buffer (50 mM Na_2_HPO_4_ pH 8.0, 0.5 M NaCl, and 8 M Urea), the beads were then washed 2 times with wash buffer containing 10 mM imidazole. After washing, the PARP7 N-terminus was eluted by incubating for 3 minutes with elution buffer (20 mM Tris pH 7.5, 100 mM NaCl, 8 M Urea, and 250 mM imidazole). Purified PARP7 N-terminus was aliquoted and flash frozen in liquid nitrogen. The antigen was sent to Pocono Rabbit Farm and Laboratory to generate the custom rabbit polyclonal antiserum against PARP7.

### Expression and purification of recombinant full-length PARP7, and determination of Km for NAD^+^ using an autoMARylation assay

Full-length, FLAG-tagged mouse PARP7 was expressed in and purified from Sf9 insect cells as previously described (Palavalli Parsons et al., 2021). For the autoMARylation assay, purified PARP7 (100 nM) was combined with NAD^+^ (0 to 2 mM) in Km Reaction Buffer (10 mM HEPES pH 7.2, 150 mM KCl, 1 mM DTT) in a reaction volume totaling 20 µL. The reaction was incubated at 25°C, 750 rpm for 10 minutes, then quenched with SDS-PAGE sample buffer. The reaction products were separated using a 4-8% PAGE-SDS gradient gel, then transferred to a nitrocellulose membrane. The membranes were then blotted for MAR and PARP7 using the MAR detection reagent and PARP7 antibody as described below. The signals were quantified with densitometry and plotted in Prism 10. The Km value was calculated with a nonlinear regression (curve fit) using the Michaelis-Menten equation.

### Generation of cell lines with stable knockdown or inducible ectopic expression

Cells were transduced with lentiviruses for Dox-inducible ectopic expression. We generated lentiviruses by transfection of the pInducer20 constructs described above, together with: (i) an expression vector for the VSV-G envelope protein (pCMV-VSV-G, Addgene 8454; RRID: Addgene_8454), (ii) an expression vector for GAG-Pol-Rev (psPAX2, Addgene 12260; RRID: Addgene_12260), and (iii) a vector to aid with translation initiation (pAdVAntage, Promega E1711) into HEK 293T cells using Lipofectamine 3000 Reagent (Invitrogen, L3000015) according to the manufacturer’s protocol. The resulting viruses were collected in the culture medium, concentrated by using a Lenti-X concentrator (Clontech, 631231), and used to infect cells. Stably transduced cells were selected with G418 sulfate (Sigma, A1720; 1 mg/mL). The cells were treated with 1 μg/mL Dox for 24 hours to induce protein expression. Inducible ectopic expression of PARP7 was confirmed by Western blotting.

### CRISPR/Cas9-mediated knockout of *Parp7* in 3T3-L1 cells

Lentiviruses were generated by transfecting the lentiCRISPR v2 vectors described above into HEK 293T cells and were used to infect 3T3-L1 cells. The infected cells were selected with 2 μg/mL puromycin (Sigma, P9620), expanded, and frozen in aliquots for future use. The efficiency of PARP7 knockout in bulk cells was verified by Western blot.

### siRNA-mediated knockdown of MART mRNAs in 3T3-L1 cells

The siRNAs for the MARTs and the control siRNA (SIC001) were purchased from Sigma. All the siRNA oligos were transfected at a final concentration of 30 nM using Lipofectamine RNAiMAX reagent (Invitrogen, 13778150) according to the manufacturer’s instructions. The cells were used for various assays at least 48 hours after siRNA transfection.

***siRNAs.*** The following siRNAs were used:

*-* PARP3 (siRNA1: SASI_Mm01_00170378, siRNA2: SASI_Mm01_00170377)

*-* PARP4 (siRNA1: SASI_Mm02_00446759, siRNA2: SASI_Mm02_00446760)

*-* PARP6 (siRNA1: SASI_Mm02_00336176, siRNA2: SASI_Mm02_00336177)

*-* PARP7 (siRNA1: SASI_Mm01_00095332, siRNA2: SASI_Mm01_00095331)

*-* PARP8 (siRNA1: SASI_Mm02_00294336, siRNA2: SASI_Mm02_00294337)

*-* PARP10 (siRNA1: SASI_Mm02_00399033, siRNA2: SASI_Mm02_00399032)

*-* PARP11 (siRNA1: SASI_Mm02_00349742, siRNA2: SASI_Mm02_00349741)

*-* PARP12 (siRNA1: SASI_Mm01_00087982, siRNA2: SASI_Mmo1_00087983)

*-* PARP14 (siRNA1: SASI_Mm01_00041267, siRNA1: SASI_Mm01_00041264)

*-* PARP16 (siRNA1: SASI_Mm01_00098868, siRNA2: SASI_Mm01_00098869)

*-* Cebpb (siRNA1: SASI_Mm01_00187563, siRNA2: SASI_Mm02_00317369)

### Lysate preparation and immunoblotting

Cells were cultured and treated as described above before the preparation of cell extracts.

***Preparation of whole cell lysates*.** For whole cell lysates at the conclusion of the treatments, the cells were washed twice with ice-cold PBS and resuspended in Lysis Buffer (20 mM Tris-HCl pH 7.5, 150 mM NaCl, 1 mM EDTA, 1 mM EGTA, 0.1% NP-40, 1% sodium deoxycholate, 0.1% SDS) containing 1 mM DTT, 250 nM ADP-HPD, 10 mM PJ-34, 1x complete protease inhibitor cocktail (Roche, 11697498001) and 1x phosphatase inhibitor cocktail (Sigma-Aldrich, P0044, P5726). The cells were resuspended in Lysis Buffer and incubated on ice for 15 minutes, vortexed for 30 seconds and then centrifuged at full speed for 15 minutes at 4°C in a microcentrifuge to remove the cell debris. Protein concentrations were measured using the Bio-Rad Protein Assay Dye Reagent (5000006) and volumes of lysates containing equal total amounts of protein were mixed with 1/4 volume of 4x SDS-PAGE Loading Solution (250 mM Tris pH 6.8, 40% glycerol, 0.04% Bromophenol Blue, 4% SDS) and boiled at 95°C for 10 minutes, except for lysates used for MAR and PAR which were heated to 65°C for 10 minutes.

***Tissue lysate preparation.*** Tissue samples were thawed on ice and were then added to HNTG buffer (50 mM HEPES pH 7.5, 150 mM NaCl, 10% Glycerol, 1% Triton X-100) plus protease inhibitor cocktail, phosphatase inhibitor cocktail and 1 mM DTT. Tissues were then minced with scissors and homogenized using the Fisher Scientific PowerGen 125 homogenizer. Lysates were then centrifuged at max speed for 20 minutes at 4°C. Protein concentrations were measured using the Bio-Rad Protein Assay Dye Reagent (5000006) and volumes of lysates containing equal total amounts of protein were mixed with 1/4 volume of 4x SDS-PAGE Loading Solution (250 mM Tris pH 6.8, 40% glycerol, 0.04% Bromophenol Blue, 4% SDS) and boiled at 95°C for 10 minutes.

***Immunoblotting.*** Protein lysates were run on SDS-PAGE gels and transferred to nitrocellulose membranes. After blocking with 3% nonfat milk in TBST, the membranes were incubated with the primary antibodies described above in TBST with 0.02% sodium azide, followed by anti-rabbit HRP-conjugated IgG (1:2000) or anti-mouse HRP-conjugated IgG (1:2000). Immunoblot signals were captured using a luminol-based enhanced chemiluminescence (ECL) HRP substrate (SuperSignal™ West Pico; Thermo Scientific, 34580) or an ultra-sensitive enhanced chemiluminescence HRP substrate (SuperSignal™ West Femto; Thermo Scientific, 34094) and a ChemiDoc imaging system (Bio-Rad).

### Immunoprecipitation of PARP7

HEK 293T cells were seeded at ∼2 × 10^6^ cells per 15 cm diameter plate and transfected at ∼60% confluence with pcDNA3 containing a cDNA encoding FLAG-tagged wild-type or mutant mouse PARP7, as described above using Lipofectamine 3000 Reagent (Invitrogen, L3000015) for 72 hours according to the manufacturer’s protocol. 3T3-L1 cells ectopically expressing PARP7 were grown to confluency and were treated with 1 μg/mL Dox to induce FLAG-tagged protein expression. Appropriate cell treatments were performed as described above. The cells were collected and whole cell extracts were prepared as described above. The resulting extracts were incubated with equilibrated anti-FLAG M2 beads (Sigma-Aldrich, A2220) for 16 hours at 4°C with gentle mixing. The beads were washed five times with gentle mixing for 10 minutes at 4°C with Immunoaffinity Purification Wash Buffer (25 mM Tris-HCl pH 7.5, 150 mM NaCl, 0.1% NP-40 and 1x complete protease inhibitor cocktail). The beads were then heated to 65°C for 10 minutes in 2x SDS-PAGE loading buffer to release the bound proteins. The immunoprecipitated material was subjected to immunoblotting as described above.

### Determining the half-life of PARP7

3T3-L1 cells were differentiated using MDI cocktail. At 0 or 8 hours of differentiation, the cells were subjected to treatment with 100 μg/mL cycloheximide (CHX) for a range of times up to 40 minutes with intervals of 5 or 10 minutes. The cells were collected and then subjected to Western blotting for PARP7 protein levels in the presence or absence of ongoing protein synthesis.

### Western blotting for PARP7 ubiquitylation and screening of E3 ubiquitin ligases

The ubiquitylation of ectopically expressed FLAG-tagged PARP7 in 3T3-L1 cells was examined using IP-Western assays as described above. Antibodies for total and K48-linked ubiquitin were used for detection. The role of various E3 ubiquitin ligases was determined by using siRNA-mediated knockdown.

***siRNAs.*** The following siRNAs were used to knockdown the E3 ligases:

*-* HUWE1 (siRNA1: SASI_Mm01_00100375, siRNA2: SASI_Mm01_00100376)

*-* DTX2 (siRNA1: SASI_Mm02_00327321, siRNA2: SASI_Mm01_00086011)

*-* UBR5 (siRNA1: SASI_Mm02_00297641, siRNA2: SASI_Mm02_00307003)

*-* RNF114 (siRNA1: SASI_Mm01_00103953, siRNA2: SASI_Mm01_00103954)

*-* TRIM28 (siRNA1: SASI_Mm01_00036667, siRNA2: SASI_Mm01_00036669)

***Primers for RT-qPCR.*** The following primers were used to confirm knockdown of the E3 ligases by RT-qPCR:

*-* HUWE1 Forward: 5’-TGAATGCTTTGGCTGCATAC-3’

*-* HUWE1 Reverse: 5’-CCCCAGGTTTAGGA TCAGATT-3’

*-* DTX2 Forward: 5’-GGCTGTGGCTTCTGGGTACAG-3’

*-* DTX2 Reverse: 5’-CCTTGTTCCCGTTGCAATACA-3’

*-* UBR5 Forward: 5’-TCCATCCATTTCGTGGTCCA-3’

*-* UBR5 Reverse: 5’-GGGTGGCTGTTCAAATTGTACTT-3’

*-* RNF114 Forward: 5’-CGGCAGATCGAGAGCATAGAG-3’

*-* RNF114 Reverse: 5’-TGGCCTTTACACCTTCCATGA-3’

*-* TRIM28 Forward: 5’-ATGGAGCAGATAGCACTGG-3’

*-* TRIM28 Reverse: 5’-GGAGAACACGCTCACATTTCC-3’

### RNA isolation and reverse transcription quantitative real-time PCR (RT-qPCR)

3T3-L1 cells or SVF cells were seeded in 6-well plates and treated as described above. The cells (or tissues) were collected, and total RNA was isolated using TRIzol Reagent (Invitrogen, 15596026) according to the manufacturer’s protocols. Total RNA was reverse transcribed using oligo (dT) primers and MMLV reverse transcriptase (Promega, M1701) to generate cDNA. The cDNA samples were subjected to qPCR using gene-specific primers, as described below. For the reverse transcription quantitative real-time PCR (RT-qPCR) analyses, “relative expression” was determined in comparison to a value of the control sample. Target gene expression was normalized to the expression of *Tbp* mRNA. The normalized value from the control sample was set to 1 and all the rest of the values were plotted against it. All experiments were performed a minimum of three times with independent biological replicates to ensure reproducibility and statistical significance.

***Primers used for RT-qPCR.*** The following gene-specific primer sequences were used:

*- Adipoq* forward: 5’-GACAAGGCCGTTCTCTTCAC-3’

*- Adipoq* reverse: 5’-CAGACTTGGTCTCCCACCTC-3’

*- Pparg* forward: 5’-TGCTGTTATGGGTGAAACTCT-3’

*- Pparg* reverse: 5’-CGCTTGATGTCAAAGGAATGC-3’

*- Fabp4* forward: 5’-AAGTGGGAGTGGGCTTTGC-3’

*- Fabp4* reverse: 5’-CCGGATGGTGACCAAATCC-3’

*- Tbp* forward: 5’-TGCTGTTGGTGATTGTTGGT-3’

*- Tbp* reverse: 5’-CTGGCTTGTGTGGGAAAGAT-3’

*- Cebpa* Forward: 5’-GAACAGCAACGAGTACCGGGTA-3’

*- Cebpa* Reverse: 5’-GCCATGGCCTTGACCAAGGAG-3’

*- Cebpb* Forward: 5’-CAAGCTGAGCGACGAGTACA-3’

*- Cebpb* Reverse: 5’-CAGCTGCTCCACCTTCTTCT-3’

*- Parp3* Forward: 5’-GCAGCACCTGCTGATAATCG-3’

*- Parp3* Reverse: 5’-TCCTCGTGGACCTGTATCCC-3’

*- Parp4* Forward: 5’-CGGCTTGAGCTGGGTAAAGA-3’

*- Parp4* Reverse: 5’-AGAGAAGTGTTGGCTTGGGG-3’

*- Parp6* Forward: 5’-TGAGGTTTTCCAGCCATCGAA-3’

*- Parp6* Reverse: 5’-GCCAGCTCGGAACTTCTTGA-3’

*- Parp7* Forward: 5’-ATGCACTGGATTTCGTGGCT-3’

*- Parp7* Reverse: 5’-AGTGGCACCGTTTCCAAGTT-3’

*- Parp8* Forward: 5’-ACACTATCCTATTGGCCTGCTG-3’

*- Parp8* Reverse: 5’-AGATGCCTAGGCTACTGCCC-3’

*- Parp10* Forward: 5’-CGGGCCTTTTATAGCACCCT-3’

*- Parp10* Reverse: 5’-GTGCCATGGTAGAGGACCTG-3’

*- Parp11* Forward: 5’-GGAGGTTCGATGGTCCGAAG-3’

*- Parp11* Reverse: 5’-TCCCACATAACGGTGCGAA-3’

*- Parp12* Forward: 5’-TTACAGGCCCAAAGAGCAT-3’

*- Parp12* Reverse: 5’-GCCACTGGTAGACTTCCCAC-3’

*- Parp14* Forward: 5’-GCCCTTGGTCTGTGTGAACT-3’

*- Parp14* Reverse: 5’-AGGGTTTCCACTGGAGAGGT-3’

*- Parp16* Forward: 5’-CAGTGCCAAGAAGGCAGAGT-3’

*- Parp16* Reverse: 5’-GGCCGGGTCAAGTACTCAA-3’

### RNA sequencing (RNA-seq) and data analysis

***RNA isolation.*** 3T3-L1 cells were grown and differentiated in a 6-well plate, as mentioned above and were transfected with siRNAs targeting *Parp7* or a universal control using the Lipofectamine 3000 Reagent (Invitrogen, L3000015). The cells were collected, and total RNA was isolated using the Direct-zol RNA MiniPrep Plus (Zymo Research, R2070) according to the manufacturer’s instructions.

***RNA-seq library preparation.*** The RNA obtained above was used to generate strand-specific RNA-seq libraries. Briefly, the libraries were prepared using the NEBNext Ultra II Directional RNA Library Prep Kit (New England Biolabs, E7760S) with the NEBNext Poly(A) mRNA magnetic Isolation Module (New England Biolabs, E7490S) for mouse SVF samples and 3T3-L1 cells. Specifically, 500 ng of total RNA was used for mRNA isolation and cDNA synthesis, 0.6 µM of adapter was used for generating the mRNA library template, and the final mRNA library was amplified with 14 x PCR cycles and sequenced on an Illumina NEXTseq2000 system by paired-end 50 bp using NEXTseq 2000 P3-100 cycle sequencing kit (Illumina, 20040559). Four biological replicates were included in final library. Publicly available data from GSE57415 was used for some analysis.

***Initial analysis of RNA-seq.*** QC analyses were performed on raw data using the FastQC tool (Andrews, 2010). The reads were then aligned to the mouse genome (mm10) using the splice junction aligner, TopHat version.2.0.12 (Kim et al., 2013). Bam files consisting of uniquely mapped reads were converted into bigWig files using BEDTools (version 2.17.0) (Quinlan and Hall, 2010) for visualization in the UCSC Genome Browser (version 2.9.4) (Perez et al., 2024). Mapped reads were counted for genomic features using featureCounts (Liao et al., 2014) followed by differential analysis of the count data with the DESeq2 package (Love et al., 2014). The significant differentially expressed genes were defined with a p-value < 0.05 and FC > 1.5 for upregulated and FC < 0.67 for downregulated.

***Data visualization.*** Heatmaps were created using Java Treeview (Saldanha, 2004) to display genes that were significantly altered in at least one experimental condition. Boxplots were generated using custom R scripts to showcase FPKM values amongst genes in different experimental conditions relative to undifferentiated or differentiated controls. Statistical significance amongst comparisons was established using Wilcoxon rank sum tests (p < 0.05).

***Gene ontology analysis.*** Gene Ontology analysis was conducted to identify enriched biological processes amongst experimental conditions. The analysis was completed using the DAVID tool (Dennis et al., 2003; Huang et al., 2009), where ontological terms were ranked based on enrichment scores.

### Chromatin immunoprecipitation (ChIP)

***Cell culture.*** 3T3-L1 cells were cultured and treated as described above in 15 cm diameter plates. The cells were treated with vehicle control DMSO, 10 µM MG132, or 400 nM RBN2397 as described above. The cells were then differentiated for 8 hours before collection as described below.

***ChIP for PARP7.*** ChIP was performed as described previously (Kim et al., 2023).

Briefly, the cells were cross-linked with 1% formaldehyde in PBS for 5 minutes at room temperature and quenched in 125 mM glycine in PBS for 5 minutes at 4°C. Cross-linked cells were then collected by centrifugation and lysed in RIPA 0 Buffer (10 mM Tris-HCl pH7.4, 1mM EDTA, 0.1% sodium deoxycholate, 0.1% SDS, 1% Triton X-100, 0.25% Sarkosyl, 1 mM DTT, and 1x complete protease inhibitor cocktail). A crude lysate was sonicated to generate chromatin fragments of ∼300 bp in length. The soluble chromatin was clarified by centrifugation, NaCl was added to a final concentration of 0.3 M. 2.5% input was removed from samples and was treated the same as the immunoprecipitation material throughout the remainder of the assay.

The samples were used in immunoprecipitation reactions with antibodies against PARP7 (in-house N-terminal antibody) or rabbit IgG (as a control) with incubation overnight at 4°C with gentle mixing. The next day, Protein A Dynabeads were washed 3 times, added to each IP sample, and were left to rotate at 4°C for 3 hours. The samples were then washed 2 times with each of the following buffers (1) RIPA 0 Buffer (10 mM Tris-HCl pH 7.4, 1 mM EDTA, 0.1% SDS, 1% Triton X-100, 0.1% sodium deoxycholate, and 1x complete protease inhibitor cocktail), (2) RIPA 0.3 Buffer (10 mM Tris-HCl pH 7.4, 1 mM EDTA, 0.3 M NaCl, 0.1% SDS, 1% Triton X-100, 0.1% sodium deoxycholate, and 1x complete protease inhibitor cocktail), (3) LiCl Wash Buffer (10 mM Tris-HCl pH 7.9, 1 mM EDTA, 250 mM LiCl, 0.5% NP-40, 0.5% sodium deoxycholate, and 1x complete protease inhibitor cocktail), and (4) 1x Tris-EDTA (TE). The immunoprecipitated genomic DNA was eluted in SDS Elution Buffer (1% SDS, 10 mM EDTA, 50 mM Tris-HCl pH 7.9) and decrosslinked at 65°C overnight. Supernatant was then collected and digested with RNase A and proteinase K to remove residual RNA and protein, respectively. ChIP DNA was then recovered using Qiagen PCR Purification Kit (28104) and eluted in DEPC- treated, nuclease-free water.

### ChIP-qPCR

The ChIPed genomic DNA was subjected to qPCR using gene-specific primers. The immunoprecipitation of genomic DNA was normalized to the input. All experiments were performed a minimum of two times with independent biological replicates.

***Primers for ChIP-qPCR.*** The following primers were used:

*- Cebpa* Forward: 5’-CTGGAAGTGGGTGACTTAGAGG-3’

*- Cebpa* Reverse: 5’-GAGTGGGGAGCATAGTGCTAG-3’

*- Pparg* Forward: 5’-GGCCAAATACGTTTATCTGGTG-3’

*- Pparg* Reverse: 5’-GTGAGGGGCGTGAACTGTA-3’

### ChIP-sequencing (ChIP-seq) and data analysis

***ChIP-seq library preparation.*** ChIP-seq libraries were generated from three biological replicates for each condition. A total of 10 ng for PARP7 ChIPed DNA, or equivalent amounts of input DNA, were used to generate libraries for sequencing. ChIP-seq libraries were generated using NEBNext Ultra II DNA Library Prep Kit (New England Biolabs, E7645) and sequenced on an Illumina NextSeq2000 system by paired-end 50 bp using NEXTseq 2000 P3-100 cycle sequencing kit (Illumina, 20040559). Three biological replicates were included in final library. Publicly available data from GSE27826 was used for some analysis (Siersbaek et al., 2011).

***Quality check and preprocessing ChIP-seq libraries.*** The raw reads were subjected to quality check using the FastQC tool (Andrews, 2010). The paired-end reads were trimmed using Cutadapt (ver. 1.9.1) (Martin, 2011) and aligned to mouse reference genome (mm10) using default parameters in Bowtie (ver. 2.2.8) (Langmead and Salzberg, 2012). The aligned reads were filtered for quality and uniquely mappable reads using Samtools (ver.0.1.19) (Li et al., 2009) and Picard (ver. 1.127, http://broadinstitute.github.io/picard/) were used for further downstream analysis. Uniquely mapped reads that met minimum ENCODE data quality standards (Landt et al., 2012) were converted to bigwig files using BEDTools (ver. 2.17.0) (Quinlan and Hall, 2010).

***Peak calling.*** Relaxed peaks were called using MACS (ver. 1.4.0) (Feng et al., 2012) with default parameters q-value = 1x10^-2^ for each sample using input condition as a control. Peaks were annotated in terms of genomic features using the ChIPseeker package in R (Yu et al., 2015).

***Motif Search.*** De novo motif search on 2000 bp region surrounding the peak summit (± 1000 bp) was performed using findMotifsGenome.pl program of HOMER motif discovery algorithm (ver. 4.9) (Heinz et al., 2010).

***Integration, analysis and visualization of ChIP-seq and RNA-seq data.*** Nearest neighboring gene for each ChIP peak was determined using GREAT (v. 3.0.0) (McLean et al., 2010) within a specific distance from the peak summit.

### CUT&RUN sequencing and data analysis

***Cell culture.*** 3T3-L1 cells were cultured and treated as described above in 15 cm diameter plates. The cells were transfected with siRNAs targeting *Parp7* or a universal control 48 hours before the addition of the differentiation cocktail. The cells were then differentiated for 24 hours before collection as described below.

***CUT&RUN library prep.*** Cells were harvested by trypsinization and counted using the BioRad TC20 automated cell counter. One-hundred thousand cells were collected and pelleted by centrifugation. CUT&RUN libraries were prepared using Epicypher kits (CUTANA ChIC/CUT&RUN Assay Kit, 14-1048, and CUTANA Library Prep Kit, 14-1001) according to manufacturer’s protocol. In short, the cells were bound to beads, permeabilized using 0.1% digitonin and incubated with 0.5 µg of antibodies against C/EBPβ or H3K27ac. Two biological replicates were collected.

***Initial analysis of CUT&RUN data.*** A quality check was performed on the raw reads using the FastQC tool. Reads were trimmed for the removal of low quality reads and adapter removal using Trimmomatic (ver. 0.39) (Bolger et al., 2014) with the following parameters: ILLUMINACLIP:Truseq3.PE.fa:2:15:4:4:true LEADING:20 TRAILING:20

SLIDINGWINDOW:4:15 MINLEN:25. The paired end reads were then aligned to the mm10 genome using bowtie2 (ver. 2.2.8) (Langmead and Salzberg, 2012). In addition to the default settings of bowtie2, the following parameters were included: –no-mixed, –no-discordant, and – dovetail. Furthermore, the paired ends were also aligned to the *E. coli* K12, MG1655 reference genome to normalize the sequencing reads to the *E. coli* spike-in DNA.

***Data normalization and peak calling.*** Normalization of sequencing reads was performed as per Epicypher’s recommendation (Meers et al., 2019). Briefly, *E.coli* spike-in reads were quantified as a percentage by examining the *E.coli* reads relative to the total read count in each condition. A normalization factor was calculated with the following formula: 1/(% *E. coli* spike-in per condition). These values were then used to generate normalized bigwig files for visualization using the deepTools bamCoverage (Ramirez et al., 2016) with the –scaleFactor option enabled. Browser tracks were generated with these bigwig files and visualized using UCSC Genome Browser. Peaks were called using MACS (ver. 2.1.0) (Zhang et al., 2008) with a *q*-value = 1x10^-2^ for each condition with the input condition being the control. Peaks were annotated in terms of genomic features using the ChIPseeker package in R (Yu et al., 2015).

***Peak annotation and clustering.*** Peaks were defined into three groups: gained, maintained, and depleted in response to the experimental conditions. The reads under the peaks were calculated for the treatment condition (T2) as well as the control (T1). Rc was calculated using the following formula: Rc = log(T1/T2). Smaller Rc values indicate binding depletion, while larger Rc values indicate enrichment. The cutoff used to define gained, maintained, and depleted peaks was the median absolute deviation (MAD) calculated for the Rc values.

***Data visualization and statistics.*** To showcase the CUT&RUN peak data as a heatmap, read densities 5 kb surrounding the gained, maintained, and depleted peaks were calculated using HOMER software (ver. 4.10.4) (Heinz et al., 2010). Heatmap visualization was completed with Java Treeview (Saldanha, 2004). Metaplots were generated using Deeptools 2.0 (Ramirez et al., 2016) to illustrate the distribution of reads near C/EBPβ binding sites with PARP7 knockdown.

### Assay for Transposase-Accessible Chromatin (ATAC) sequencing and data analysis

**Cell culture.** 3T3-L1 cells were cultured and treated as described above in 15 cm diameter plates. The cells were transfected with siRNAs targeting *Parp7* mRNA or a universal control 24 hours before the addition of the differentiation cocktail. The cells were then differentiated for 24 hours before collection as described below. Cells were harvested by trypsinization and counted using the BioRad TC20 automated cell counter. Fifty-thousand cells were collected and pelleted by centrifugation.

***ATAC- Seq sample preparation.*** Samples were then resuspended in buffer containing 10 mM Tris-HCl pH 7.4, 10 mM NaCl, 3 mM MgCl_2_, 0.1% NP-40, 0.1% Tween-20, and 0.01% digitonin, and incubated on ice for 3 minutes. Nuclei were then washed twice with buffer containing 10 mM Tris-HCl pH 7.4, 10 mM NaCl, 3 mM MgCl_2_, 0.1% Tween-20. Nuclei were pelleted by spinning at 500 x *g* for 10 minutes at 4°C in a fixed angle centrifuge. Pellets were then resuspended in transposition mixture containing 20 mM Tris-HCl pH 7.6, 10 mM MgCl_2_, 20% DMF, 0.01% Digitonin, 0.1% Tween-20, and 100 nM recombinant Tn5 transposase. After pipetting up and down 6 times, mixture was incubated at 37°C for 30 minutes with 1000 RPM mixing using the Eppendorf Thermomixer C. The resulting tagmented DNA was purified using Qiagen PCR CleanUp columns and was eluted with elution buffer. Samples were then combined with 2X NEBNext High-Fidelity Master Mix (NEB, M0541L), Nextera i7 and i5 indexed primers, and amplified by PCR: (1) 72°C 5 minutes, (2) 98°C 30 seconds, (3) 98°C 10 seconds,

1. (4) 63°C 10S seconds. PCR steps 3-4 were repeated 8 times. Adaptor contamination was removed from libraries using 1.3x volume AMPure beads and eluted in 10 mM Tris-HCl (pH 8). Two biological replicates were included.

***Quality check and preprocessing ATAC-seq libraries.*** To check the quality of ATAC-seq libraries the reads were assessed using the FastQC tool (Andrews, 2010). The raw reads were then aligned to mouse genome (mm10) using BWAKit ver.0.7.15 (Li and Durbin, 2009). The aligned reads were then subjected to a quality score check, assessed for unique alignments using Picard tools (ver. 2.10.3, http://broadinstitute.github.io/picard/) and only the reads with a score over 10 were used for further analysis. The uniquely mapped reads were normalized for read depth across all samples and converted to bigWig files using the writeWiggle function from the groHMM package in R (Chae et al., 2015) and visualized on the UCSC genome browser.

### Mouse studies

All animal experiments were performed in compliance with the Institutional Animal Care and Use Committee (IACUC) at the UT Southwestern Medical Center. The *Parp7* tm1a [“knockout-first” allele; (Skarnes et al., 2011)] mouse strain used for this research project, C57BL/6N-*A^tm1Brd^ Tiparp^tm1a(EUCOMM)Wtsi^*/TcpMmucd (RRID:MMRRC_050063-UCD) was obtained from the Mutant Mouse Resource and Research Center (MMRRC) at the University of California at Davis, an NIH-funded strain repository, and was donated to the MMRRC by The KOMP Repository, University of California, Davis; Originating from Colin McKerlie, The Toronto Centre for Phenogenomics. The mice were re-derived from cryopreserved sperm.

### Primers for genotyping the mice

*-* PARP7_tm1C Forward: 5’-TTGAATCAGCACTACTGGCCTC-3’

*-* PARP7_tm1C Reverse: 5’-GACAGCCTTCGTAGTTGGTCA-3’

*-* PARP7_floxed Forward: 5’-ACAGAGTTCTGAAAAGAGGATTTGCC-3’

*-* PARP7_floxed Reverse: 5’-GAGATGGCGCAACGCAATTAAT-3’

*-* Wild-type Forward: 5’-CAGCACTACTGGCCTCATCTGG-3’-3’

*-* Wild-type Reverse: 5’-CTCGAAGTCAATCAGTGAGTCAGC-3’

***High-fat diet (HFD).*** Eight-week-old male PARP7 knockdown (*Parp7^tm1a/tm1a^*) and control (*Parp7^WT/WT^*) mice were switched from standard chow diet to chow diet containing 60% fat for eight weeks. Body weights were monitored weekly. Body composition was measured using an EchoMRI-100 Body Composition Analyzer (UT Southwestern Metabolic Phenotyping Core). At the end of the experiment tissues were collected and flash frozen in liquid nitrogen.

***Mammary pad involution.*** Female mice were crossed with male mice and separated into their own cage before giving birth to pups. Two mating strategies were set up: (1) PARP7 knockdown (*Parp7^tm1a/tm1a^*) females were crossed to PARP7 control (*Parp7^WT/WT^*) males, (2) and the PARP7 control (*Parp7^WT/WT^*) females were crossed to PARP7 knockdown (*Parp7^tm1a/tm1a^*) males. This ensured all pups born were heterozygous (*Parp7^WT/tm1a^*). Mothers were left with their litter for 12 days for mammary fat dedifferentiation to occur as previously described (Wang et al., 2018). After 12 days of feeding, pups were removed from the mother to allow for involution to occur for 3 days. Time points were collected at weaning (day 0) and after 3 days of involution (day 3). At the point of collection, tissues to be used for Western blotting were flash frozen in liquid nitrogen or fixed as described below.

### Immunofluorescent staining of tissue

Tissues were fixed in 10% formalin for 24 hours and stored in 100% ethanol. Paraffin embedding, slicing and mounting was done by the UTSW Histopathology Core. Slides were baked on 65°C heat block for 30 minutes and were cooled at room temperature for 15 minutes. For hydration slides were rinsed 3 times 5 minutes each in xylene to remove residual paraffin, rinsed 3 times in 100% ethanol for 3 minutes each, then the following washes were preformed twice for 3 minutes each in 95% ethanol, 70% ethanol, 50% ethanol, deionized water. After washes slides were incubated in PBS for 3 minutes. Antigen retrieval was performed by incubating slides in boiling antigen unmasking solution (Vector Labs, H-3300-250) containing 0.05% Tween20 for 12 minutes and allowing them to stay in the solution for 18 more minutes. Slides were then cooled at room temperature for 30 minutes and were rinsed in PBS for 3 minutes. Slides were blocked with 10% normal goat serum in PBST for 50 minutes at room temperature. Primary antibody was added in a 1:1000 dilution in PBST and incubated overnight at 4°C in a humidified chamber. After primary antibody incubation slides were removed from chamber and were rinsed 5 times in PBST for 10 minutes at room temperature. Secondary antibody was diluted in 10% normal goat serum in PBST and was incubated at room temperature for 60 minutes in a humidified chamber, slides were then rinsed 5 times in PBST for 10 minutes each. Excess moisture was removed from slides and mounting medium containing DAPI was applied, a coverslip was then added and sealed with nail polish. Images were acquired using a Keyence BZ-X810 fluorescence microscope.

### Metabolite extraction and analysis for polar metabolites

Mammary fat pad tissue (10 to 20 mg) and breast milk (5 μL) were collected. The tissue was homogenized manually with a rubber Dounce homogenizer in ice-cold acetonitrile:water (80:20) solution. The samples were flash frozen three times in liquid nitrogen and then centrifuged at 14,000 x *g* for 10 minutes at 4°C. The protein concentrations of the supernatants were determined by BCA assays, normalized to 70 μg/mL, and placed in LC–MS vials. Metabolite analysis used a Vanquish UHPLC coupled to a Thermo Scientific QExactive HF-X hybrid quadrupole orbitrap high-resolution mass spectrometer (HRMS) as described previously (Solmonson et al., 2022). LC-MS/MS data were collected using Thermo Scientific XCalibur 4.1.50 and the data were analyzed using Thermo Scientific Trace Finder v5.1.

### Lipid extractions and analysis by LC/MS

Mammary fat pad tissue (10 to 20 mg) and breast milk (5 μL) were transferred to a 1:1:2 ratio of methanol, water, and chloroform and vortexed thoroughly. Extracts were centrifuged at 10,000 x *g* for 10 minutes to separate the extraction solvent into two layers. The bottom chloroform layer was removed, transferred to a clean glass tube, and dried under a stream of purified nitrogen gas. Samples were taken up in a 65/35 mixture of isopropanol and methanol and placed into vials for LC/MS analysis.

Separation of lipids was performed using a protocol adapted from the literature (Shin et al., 2020). Briefly, we used a Thermo Scientific Accucore C18 column (2.1 mm x 150 mm) heated to 60°C and a binary solvent gradient. Mobile phase A was a 50/50 mixture of water and acetonitrile and mobile phase B was a 90/8/2 mixture of isopropanol, acetonitrile, and water. Each mobile phase contained 5 mM ammonium formate with 0.1% formic acid, which are necessary for lipid ionization. The flow rate of the mobile phase was 0.17 mL/min with the following gradient: 0-1 minutes, 10% B; 1-7 minutes, linear ramp to 60% B; 7-17 minutes, linear ramp to 70% B, 17-22 minutes, linear ramp to 100% B; 22-33 minutes, 100%B; 33-33.1 minutes, linear ramp to 10% B; 33.1-40 minutes, re-equilibration to 10% B.

Mass spectrometry analysis of lipids was performed on a Thermo Scientific Fusion Lumos 1M tribrid mass spectrometer using an acquisition method modified from the literature (Rampler et al., 2018). To accurately assign the fatty acyl composition of each lipid class reported, we injected samples twice, once in positive polarity and once in negative polarity. Each polarity uses a combination of high resolution MS1 and MS2 data from the orbitrap and low resolving power MS3 scans in the ion trap for fatty acyl assignment. Our method performs these scan events by using a combination of neutral loss and fragment ion triggers to perform successive MSn analyses as outlined in the reference. We used an inclusion list for the top 20 to 30 lipids identified in human plasma according to the NIST lipid database (Bowden et al., 2017). Fatty acyl compositions of glycerophospholipids, lysophospholipids, sphingomyelin, and fatty acids were identified with the negative polarity injection, while fatty acyl compositions of glycerolipids and ceramides were identified in the positive polarity injection. In addition to lipids on the inclusion list, we also included a top 20 data dependent acquisition schema to identify lipids that fall outside of our inclusion list. Analysis of the lipid data was performed using CompoundDiscoverer 3.3 software (Thermo Fisher Scientific) searching an in silico database of lipid species. Each matching spectrum was reviewed for accuracy at the MS1, MS2, and MS3 levels to confirm a positive identification. Separate standards for 15S-HETE and 13- HODE were run to provide accurate identification of those peaks.

### Statistical analysis of LC/MS data

Relative metabolite abundance was determined by integrating the chromatographic peak area of the precursor ion searched within a 5 ppm tolerance and then normalized to total ion count (TIC). If the peak for a metabolite was not observed in a given sample, a value of 1/5 the minimum observed amount to avoid zero values for downstream analysis. Prior to analyzing statistical significance of differences among groups, we tested whether data were normally distributed and whether variance was similar among groups. To test for normality, we performed the Shapiro–Wilk tests (significantly altered normality was considered if p < 0.05). To test whether variability significantly differed among groups we performed *F*-tests (Unequal variability was considered if p<0.05). When the data significantly deviated from normality or variability significantly differed among conditions, we log_2_-transformed the data and tested again for normality and variability. If the transformed data no longer significantly deviated from normality and equal variability, we performed unpaired, two tailed Student’s t-tests on the transformed data. If log_2_-transformation was not possible or the transformed data still significantly deviated from normality or equal variability, we performed non-parametric tests on the non-transformed data. If the transformed data was distributed normality but significantly deviated from equal variability, we performed parametric, unpaired, two tailed Student’s t-test with a Welch’s correction.

### Additional analyses of metabolomics data

We performed additional analyses of the metabolomics data using MetaboAnalyst software (https://www.metaboanalyst.ca) (Pang et al., 2024; Xia et al., 2009), including a derivative of a random forest analysis to determine which metabolites distinguish the metabolic profiles from the *Parp7^WT/WT^* and *Parp7^tm1a/tm1a^*samples. The proteomics dataset was imported into MetaboAnalyst for statistical analysis. To reduce batch effects, one-factor normalization strategy was used. The metabolite abundance values were normalized using median normalization, followed by a log transformation (base10) to stabilize variance and reduce skewness. We used mean centering to further adjust the data and perform downstream analysis. After normalization Random Forest was utilized for classification and feature selection.

### Quantification and statistical analyses

All sequencing-based genomic experiments were performed a minimum of two times with independent biological samples. Statistical analyses for the genomic experiments were performed using standard genomic statistical tests as described above. All gene-specific qPCR-based experiments were performed a minimum of three times with independent biological samples. All Western blotting experiments, except for those accompanying genomic experiments which were performed in the same biological replicates as the sequencing, with quantification were performed a minimum of three times with independent biological samples and analyzed by Image Lab 6.0. Statistical analyses were performed using GraphPad Prism 10. All tests and p values are provided in the corresponding figures or figure legends. In all figures, the p values are shown as: *, p < 0.0332; **, p < 0.0021; ***, p < 0.0002, ****, p < 0.0001, or as other values as specified.

## Acknowledgements

The authors would like to thank the following: (1) members of the Kraus lab for continued input and feedback on this project, specifically H.B. Kim for input on the genomic analysis and K. Pekhale for input on antibody affinity purification; (2) the UT Southwestern Metabolic Phenotyping Core, supported by NIH P30 DK127984 awarded to the Nutrition Obesity Research Center (NORC); (3) the UT Southwestern Histopathology Core; and (4) Philipp Scherer and his lab members for input and advice regarding *in vivo* experiments.

## Funding

This work was supported by a grant from the NIH/NIDDK (R01 DK069710) and funds from the Cecil H. and Ida Green Center for Reproductive Biology Sciences Endowment to W.L.K., and predoctoral fellowship from the American Heart Association to M.S.S. In addition, this work was supported by the NIH/NIDDK (P30 DK127984). The content is solely the responsibility of the authors and does not necessarily represent the official views of the National Institutes of Health.

## Author Contributions

M.S.S. and W.L.K. conceived and developed this project, designed the experiments, and oversaw their execution. M.S.S. preformed most of the experiments and analyzed the data with assistance as follows: Y.J.K. performed the CUT&RUN assays and assisted with the library preparation for ChIP-seq and RNA-seq; Y.K. performed all assays related to PARP7 ubiquitylation and E3 ligases, S.K. analyzed the RNA-seq, ChIP-seq, CUT&RUN, and ATAC-seq data with assistance from T.N.; S.P.C. assisted with all aspects of the mouse work, including the breeding experiments, mammary gland involution experiments, and other assays with tissue samples, M.D. performed the PARP7 Km analyses; J.Z. and W.L.K. made the PARP7 antibody; D.H. assisted with data interpretation and writing, and performed cellular and biochemical analyses of C/EBPβ with assistance from Y.K.; T.P.M. and A.S. performed the metabolomics assays and initial data analyses; and CVC performed the final metabolomics data analyses and visualization. M.S.S. prepared the initial drafts of figures and text, which were edited and finalized by W.L.K. and C.V.C. with input from other authors. C.V.C. provided guidance and intellectual support for the mouse studies. W.L.K. secured funding to support this project, and provided intellectual support and overall leadership for the work.

## Competing Interests

W.L.K. is a holder of U.S. patent number 9,599,606, covering the ADP-ribose detection reagents used here, which have been licensed to and are sold by EMD Millipore.

## Data and Materials

The new genomic data sets generated for this study can be accessed from the NCBI’s Gene Expression Omnibus (GEO) repository (http://www.ncbi.nlm.nih.gov/geo) using the following accession numbers: Super series containing all of the data sets: GSE282161. Individual data sets: GSE282094 (RNA-seq), GSE282093 (PARP7 ChIP-seq), GSE282095 (C/EBPβ and H3K27ac CUT&RUN), GSE282092 (ATAC-seq). All of the metabolomics data sets (raw and processed) can be obtained as Excel spreadsheets from the corresponding author.

## Supplemental Figures Legends

Supplemental Figure S1. An siRNA screen of MARTs implicates PARP7 in adipogenesis.

(A) Bar graphs showing the siRNA-mediated knockdown of individual MART mRNAs in 3T3- L1 cells, as assayed by RT-qPCR. Two siRNAs for each MART were used. Each bar represents the mean + SEM; n = 3. Bars marked with asterisks are significantly different from control; Student’s t-test; * = p < 0.0332, ** = p < 0.0021, *** = p < 0.0002, **** = p < 0.0001.

(B) Bar graphs showing expression of the adipogenic marker genes *Adipoq* and *Fabp4* in 3T3-L1 cells after siRNA-mediated knockdown of the indicated MARTs, followed by differentiation using the MDI cocktail, as assayed by RT-qPCR at day 3 of differentiation. Each bar represents the mean + SEM; n = 3. Bars marked with asterisks are significantly different from control; Student’s t-test; * = p < 0.0332, ** = p < 0.0021, **** = p < 0.0001.

(C) *(Top)* Lipid accumulation in 3T3-L1 cells after siRNA-mediated knockdown of the indicated MARTs, followed by adipocyte differentiation using the MDI cocktail, assayed by Oil Red-O staining at day 8 of adipocyte differentiation. *(Bottom)* Bar graphs showing quantification of the Oil Red-O staining from the panel above. Each bar represents the mean + SEM; n = 3. Bars marked with asterisks are significantly different from control; Student’s t-test; * = p < 0.0332, ** = p < 0.0021, *** = p < 0.0002, **** = p < 0.0001.

Supplemental Figure S2. Validation of a PARP7 antiserum.

(A) Schematic representation of the PARP7 protein showing the 324 amino acid amino-terminus that was used as an antigen to generate the polyclonal antibody in rabbits.

(B) Western blot showing the specificity of the PARP7 antiserum using 3T3-L1 extracts with control or *Parp7* siRNA-mediated knockdown. β-tubulin was used as a loading control.

(C) Immunofluorescent staining of 3T3-L1 cells showing nuclear PARP7 staining with control or *Parp7* siRNA-mediated knockdown. DNA was stained with DAPI. *(Top panels)* Scale bar = 25 µm. *(Bottom panels)* Lower magnification of the merged image above. Scale bar = 100 µm.

(D) Western blot using the PARP7 antiserum showing stabilization of PARP7 in 3T3-L1 cells treated with 10 µM MG132 or 400 nM RBN2397 for 8 hours. β-tubulin was used as a loading control.

Supplemental Figure S3. Genetic depletion of PARP7 in 3T3-L1 cells and primary preadipocytes inhibits adipogenesis; siRNA-mediated depletion of DTX2 and RNF114 stabilizes PARP7.

(A) Western blot showing decreased PARP7 protein in 3T3-L1 cells with CRISPR/Cas9- mediated knockout of *Parp7*. β-tubulin was used as a loading control.

(B and C) Bar graphs showing expression of the adipogenic marker genes *Fabp4* and *Adipoq* in 3T3-L1 cells subjected to CRISPR/Cas9-mediated knockout of *Parp7* and differentiated with MDI cocktail. The expression of the adipogenic markers genes *Fabp4* (I) and *Adipoq* (J) was assayed by RT-qPCR on day 3 of differentiation. Target gene expression was normalized to the expression of *Tbp* mRNA. Each bar represents the mean + SEM; n = 3. Bars marked with asterisks are significantly different from control; ANOVA; *** = p < 0.0002.

(D) Microscopy images of BODIPY-stained lipids in 3T3-L1 cells subjected to CRISPR/Cas9- mediated knockout of *Parp7*, differentiated with MDI cocktail, and assayed at day 7 of differentiation. DNA was stained with DAPI. Scale bars = 50 µm.

(E through J) Expression of *Parp7* and adipogenic marker genes in primary preadipocytes from SVF after siRNA-mediated knockdown of *Parp7*, followed by differentiation using the MDI cocktail, as assayed by RT-qPCR at day 3 of differentiation. Bar graphs showing (E) the extent of *Parp7* knockdown and the effects of *Parp7* knockdown on the expression of adipogenic marker genes (F) *Cebpb*, (G) *Cebpa*, (H) *Pparg*, (I) *Fabp4*, and (J) *Adipoq*, as assayed by RT-qPCR at day 3 of differentiation. Target gene expression was normalized to the expression of *Tbp* mRNA. Each bar represents the mean + SEM; n = 3. Bars marked with asterisks are significantly different from control; ANOVA; * = p < 0.0332, ** = p < 0.0021, *** = p < 0.0002, **** = p < 0.0001.

(K) Bar graph showing quantification of Oil Red-O staining at day 8 of differentiation in primary preadipocytes from SVF subjected to siRNA-mediated knockdown of *Parp7*. Each bar represents the mean + SEM; n = 3. Bars marked with asterisks are significantly different from control; Student’s t-test; *** = p < 0.0002, **** = p < 0.0001.

(L through P) Confirmation of the knockdown of mRNAs encoding E3 ligases. 3T3-L1 cells were subjected to knockdown with control siRNA or siRNAs targeting *Huwe1*, *Dtx2*, *Ubr5*, *Rnf114*, and *Trim28*. Bar graphs show expression of the mRNAs encoding the E3 ligases as assayed by RT-qPCR. Target gene expression was normalized to the expression of *Tbp* mRNA. Each bar represents the mean + SEM; n = 3. Bars marked with asterisks are significantly different from control; Student’s t-test; ** = p < 0.01, *** = p < 0.001, **** = p < 0.0001.

(Q and R) Confirmation of the knockdown of E3 ligases DTX2 and RNF114. 3T3-L1 cells were subjected to knockdown with control or siRNAs targeting (Q) *Dtx2* or (R) *Rnf114*. Western blots showing the levels of (Q) DTX2 or (R) RNF114. β-tubulin was used as a loading control.

Supplemental Figure S4. PARP7 depletion regulates proadipogenic gene expression.

(A) Western blot showing decreased PARP7 protein in 3T3-L1 cells with siRNA-mediated knockdown of PARP7 at day 2 of differentiation using MDI cocktail. β-tubulin was used as a loading control.

(B) Box plots showing the expression (FPKM) of *Parp7 (left)* and *Cebpb (right)* mRNAs in 3T3- L1 cells subjected to control or *Parp7* siRNA-mediated knockdown and differentiation with MDI cocktail, as assayed by RNA-seq at day 0 and 2 of differentiation. Bars marked with different letters are significantly different from each other. Unpaired t-test (two-tailed), all p-values at least p < 0.02 for *Parp7* mRNA and p < 0.05 for *Cebpb* mRNA.

(C) Box plots showing the log_2_(fold change) of upregulated *(left)* and downregulated *(right)* genes in 3T3-L1 cells subjected to control or *Parp7* siRNA-mediated knockdown and differentiation with MDI cocktail, as assayed by RNA-seq at day 0 or day 2 of differentiation. Bars marked with different letters are significantly different from each other. Wilcoxon rank sum test, p < 2.2 x 10^-16^.

(D) Representative RNA-seq browser track showing reduced expression of the *Adipoq* gene in response to siRNA-mediated knockdown of *Parp7* at day 0 and 2 of differentiation in 3T3-L1 cells.

(E) Gene ontology terms from genes upregulated at 2 days of differentiation with *Parp7* knockdown in 3T3-L1 cells.

Supplemental Figure S5. PARP7 binds to chromatin genome-wide.

(A) Bar graphs showing relative enrichment of PARP7 *(left)* and C/EBPβ *(right)* binding at the promoters of target genes *Pparg* and *Cebpb*, as assayed by ChIP-qPCR, with or without CRISPR/Cas9-mediated knockout of *Parp7* in 3T3-L1 cells differentiated with MDI cocktail. Each bar represents the mean + range; n = 2.

(B and C) Browser tracks of genomic data at the (B) *Pparg* gene and (C) *Cebpa* gene. 3T3-L1 cells were subjected to control or *Parp7* knockdown, followed by PARP7 ChIP-seq or C/EBPβ CUT&RUN. PARP7 ChIP-seq was performed after treatment with 400 nM RBN2397 to stabilize the PARP7 protein.

(D) Bar graph showing the percent of PARP7 peaks found at promoter or enhancer regions in 3T3-L1 cells revealed by PARP7 ChIP-seq, which was performed after treatment with 400 nM RBN2397 to stabilize the PARP7 protein.

(E) Pie charts showing the distribution of PARP7 peaks found at different genomic regions in 3T3-L1 cells revealed by PARP7 ChIP-seq, which was performed after treatment with 400 nM RBN2397 to stabilize the PARP7 protein.

(F-H) Analysis of PARP7 ChIP-seq peak frequency in (F) control, (G) 400 nM RNB2397-, or (H) 10 µM MG132-treated 3T3-L1 cells, showing the distance of the peaks to the TSS of the nearest PARP7-regulated gene.

(I and J) Motif enrichment at significant PARP7 ChIP-seq peaks in 3T3-L1 cells for the (I) control and (J) 400 nM RBN2397 treatment groups.

Supplemental Figure S6. Validation of C/EBP(J CUT&RUN and effects of PARP7 depletion.

(A and B) Browser tracks showing C/EBPβ CUT&RUN (this study) compared to C/EBPβ ChIP-seq (Siersbaek et al., 2014) in 3T3-L1 cells at C/EBPβ target genes: (A) *Cebpa* and (B) *Pparg*.

(C) Bar graphs showing percent of C/EBPβ CUT&RUN peaks found at promoter or enhancer regions in 3T3-L1 cells. The data are separated by all C/EBPβ peaks *(left)*, as well as those depleted *(middle)* or maintained *(right)* in response to siRNA-mediated knockdown of *Parp7*.

(D and E) Enrichment of H3K27ac CUT&RUN reads at C/EBPβ peaks depleted upon siRNA-mediated knockdown of *Parp7* in 3T3-L1 cells. (D) Metaplots and (E) boxplots. For the boxplots, bars marked with different letters are significantly different from each other. Wilcox rank sum test, p < 2.2 x 10^-16^.

(F and G) Enrichment of ATAC-seq reads at C/EBPβ peaks depleted upon siRNA-mediated knockdown of *Parp7* in 3T3-L1 cells. (F) Metaplots and (G) boxplots. For the boxplots, bars marked with different letters are significantly different from each other. Wilcox rank sum test, the p-value for *a* is < 0.0002 and for the others is p < 0.028.

(H) Boxplots showing enrichment of ATAC-seq reads at all PARP7 peaks in 3T3-L1 cells. Bars marked with different letters are significantly different from each other. Wilcox rank sum test, the p-value for *a* is < 0.001 and for the others is p < 0.193.

(I) Bar graph showing the number of PARP7 peaks from PARP7 ChIP-seq within 20 kb of 577 C/EBPβ-regulated genes from RNA-seq in 3T3-L1 cells compared to 577 randomly selected genes.

Supplemental Figure S7. Genetic depletion of PARP7 reduces adipogenesis in vivo.

(A and B) Bar graphs showing quantification of the weight of *Parp7^WT/WT^* or *Parp7^tm1a/tm1a^* mice

(A) at 0 weeks or (B) after 8 weeks on a 60% high fat diet. Each bar represents the mean + SEM; n = 12. Bars marked with asterisks are significantly different from the control; Student’s t-test; ns = not significant, * = p < 0.0332.

(C) Bar graph showing quantification of the amount of fat, determined by MRI, in *Parp7^WT/WT^* or

*Parp7^tm1a/tm1a^* mice at 0 weeks, before starting a 60% high fat diet. Each bar represents the mean

+ SEM; n = 12. Bars marked with asterisks are significantly different from the control; Student’s t-test; * = p < 0.0332.

(D) Levels of PARP7 in *Parp7^WT/WT^*and *Parp7^tm1a/tm1a^* primary preadipocytes from SVF. Western blots showing increased PARP7 protein levels at 8 hours of differentiation using MDI cocktail in *Parp7*^WT/WT^, but not in *Parp7^tm1a/tm1a^*, SVF cells. β-tubulin was used as a loading control.

(E) Western blot showing stabilization of PARP7 in *Parp7^WT/WT^*, but not in *Parp7^tm1a/tm1a^*, primary preadipocytes from SVF treated with 400 nM RBN2397 compared to control and differentiated for 8 hours using MDI cocktail. β-tubulin was used as a loading control.

(F through H) Genetic depletion of *Parp7* in primary preadipocytes from SVF inhibits adipogenesis. Preadipocytes from *Parp7^WT/WT^* or *Parp7^tm1a/tm1a^*mice were induced to differentiate using MDI cocktail. Bar graphs showing expression of (F) *Parp7* mRNA, and adipogenic marker genes (G) *Adipoq* and (H) *Fabp4*, as assayed by RT-qPCR on day 4 of differentiation. Target gene expression was normalized to the expression of *Tbp* mRNA. Each bar represents the mean + SEM; n = 3. Bars marked with asterisks are significantly different from the control; Student’s t-test; * = p < 0.0332, ** = p < 0.0021, **** = p < 0.0001.

(I) Microscopy images of BODIPY-stained lipids in *Parp7^WT/WT^* or *Parp7^tm1a/tm1a^*primary preadipocytes from SVF differentiated for 7 days using MDI cocktail. DNA was stained with DAPI. Scale bars represent 250 µm.

(J) Representative gross anatomical images of mammary glands from female *Parp7^WT/WT^* or

*Parp7^tm1a/tm1a^* mice at day 0 or day 3 of mammary gland involution.

(K and L) Representative microscopy images of (K) H&E-stained and (L) perilipin-stained mammary fat pad tissue from female *Parp7^WT/WT^*or *Parp7^tm1a/tm1a^* mice at day 0 or day 3 of mammary gland involution. DNA was stained with DAPI. Scale bars represent (K) 100 µm and (L) 20 µm.

Supplemental Figure S8. Metabolomic analyses of mammary fat pad tissue and milk from

*Parp7^WT/WT^* and *Parp7^tm1a/tm1a^* mice.

(A) Breeding and delivery table from *Parp7^WT/tm1a^* x *Parp7^WT/tm1a^* crosses. Expected and actual results are listed for the parameters indicated.

(B) Lipidomics analysis in milk from *Parp7^WT/WT^* and *Parp7^tm1a/tm1a^* mice (n = 4 mice for each group). Samples were collected at the time of weaning (day 12 post-partum, day 0 of involution). The samples were subjected to mass spectrometry-based lipidomics. The top 50 altered lipids are shown, but none were significant.

(C) Box plots showing changes in 15S-HETE *(left)* and 13-HODE *(right)* levels in milk from *Parp7^WT/WT^* and *Parp7^tm1a/tm1a^* mice (n = 4 mice for each group). Samples were collected at the time of weaning (day 12 post-partum, day 0 of involution). The samples were subjected to mass spectrometry-based metabolomics. Unpaired t-test; n.s. = not significant.

(D and E) Metabolomics analysis in mammary fat pads and milk from *Parp7^WT/WT^* and

*Parp7^tm1a/tm1a^* mice (mammary fat pads, n = 4 mice for *WT*, n = 5 mice for *tm1a*; milk, n = 4 mice for each group). Samples were collected at the time of weaning (day 12 post-partum, day 0 of involution). The samples were subjected to mass spectrometry-based metabolomics. Heat maps showing changes in metabolites in (D) mammary fat pads and (E) milk. Eight of the 50 metabolites shown for mammary tissue were significant. All 64 of the metabolites shown for milk were significant.

(F and G) Classification and feature selection using MetaboAnalyst. (F) Enrichment analysis results and (G) Classification and feature selection performed using Random Forest (variables of importance) generated using the web-based MetaboAnalyst tool.

