## Supplemental Figures S1-S8 for "NAD^+^ Sensing by PARP7 Regulates the C/EBPβ-Dependent Transcription Program in Adipose Tissue In Vivo"

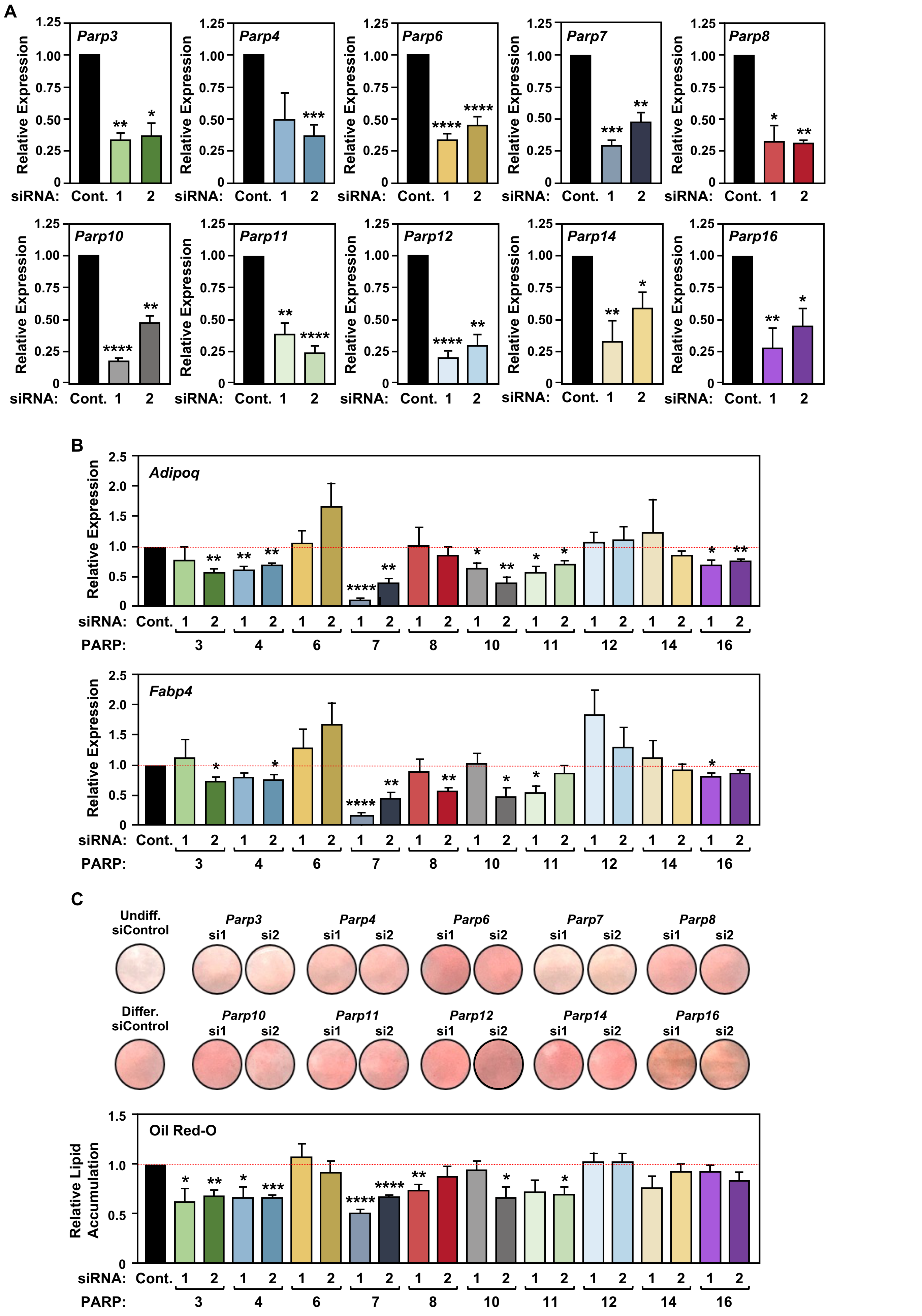


**Suppl. Fig. S1 – Stokes *et al.* (2025)**

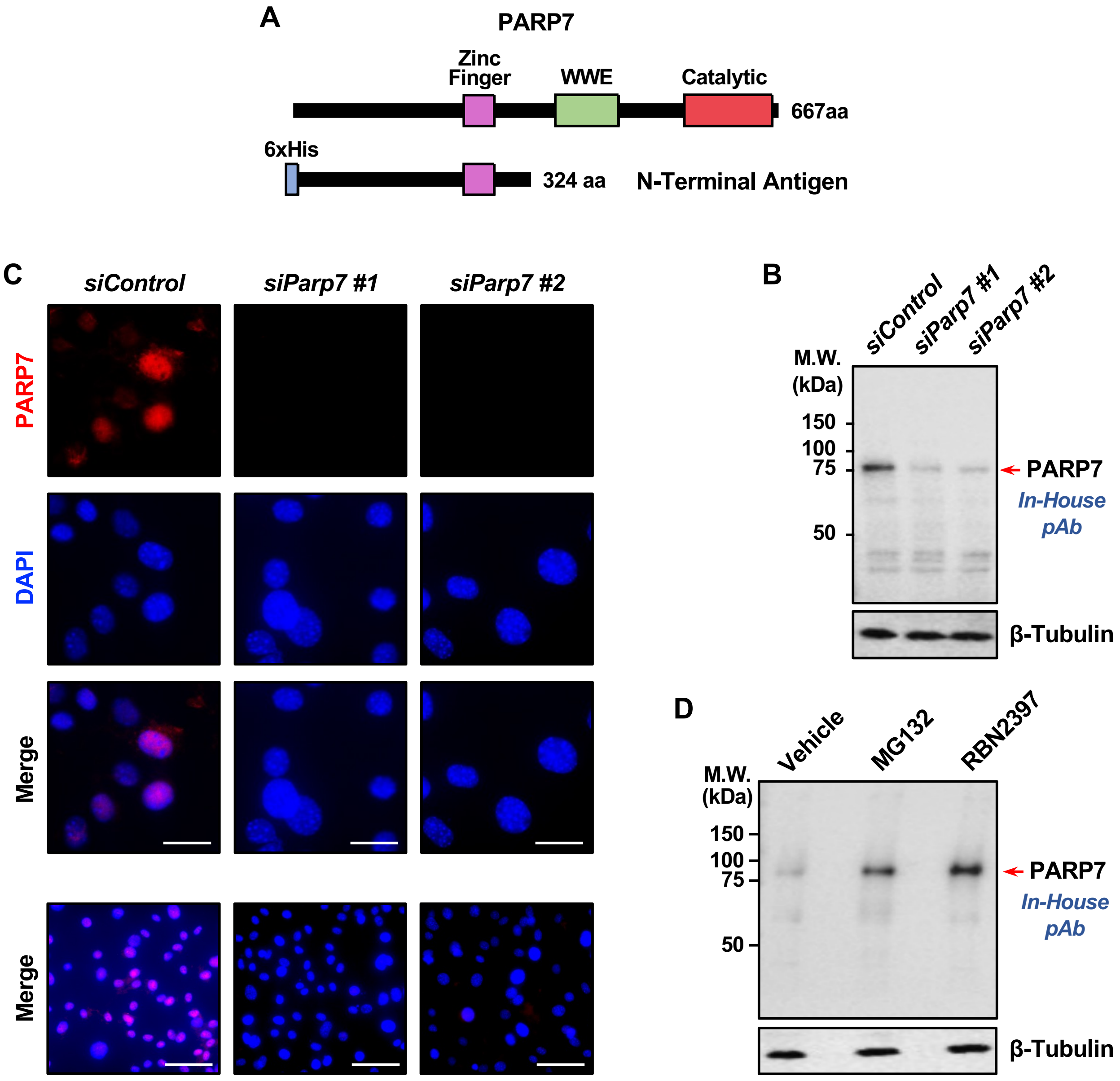


Suppl. Fig. S2 – Stokes *et al.* (2025)

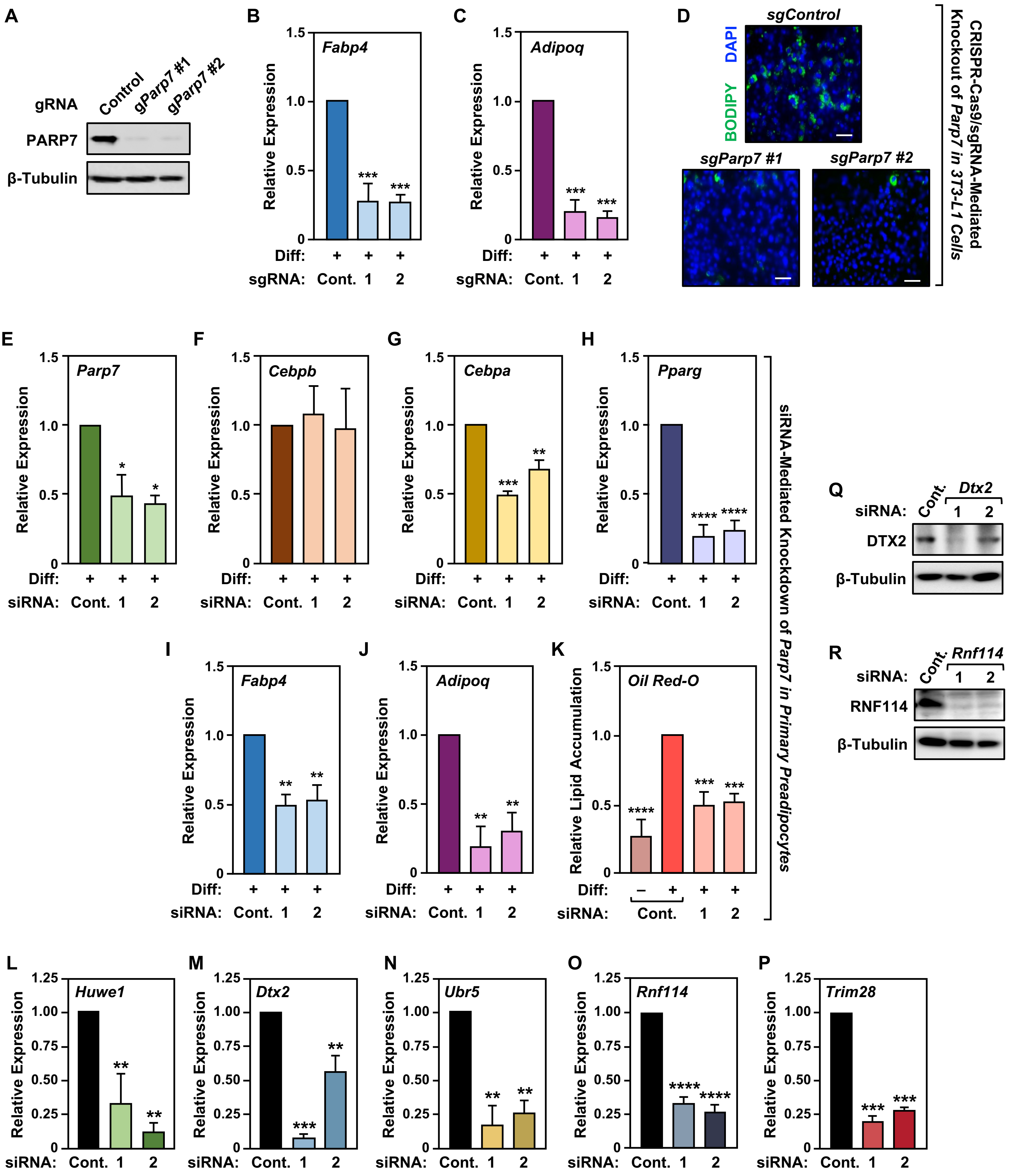


Suppl. Fig. S3 – Stokes *et al.* (2025)

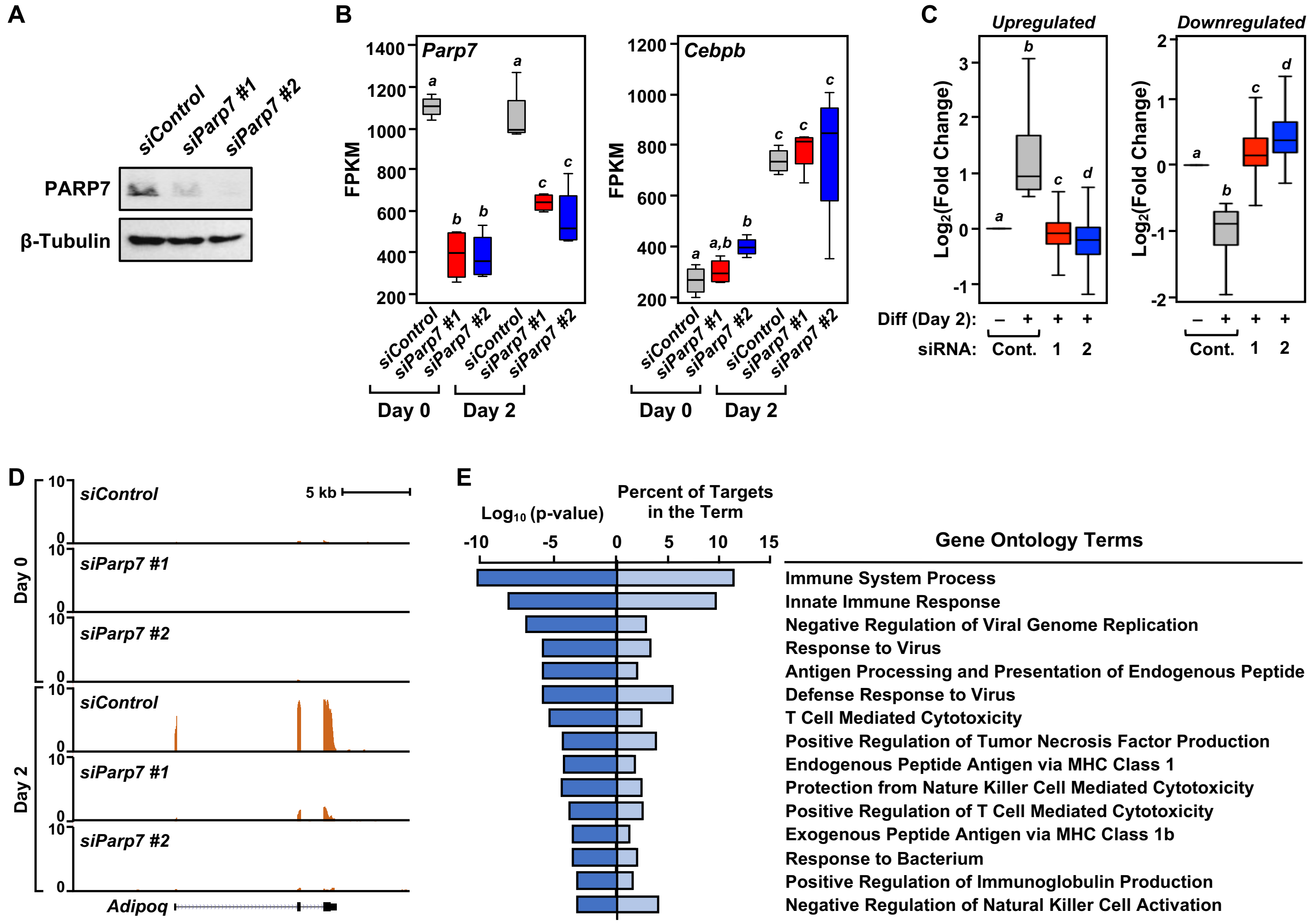


Suppl. Fig. S4 – Stokes *et al.* (2025)

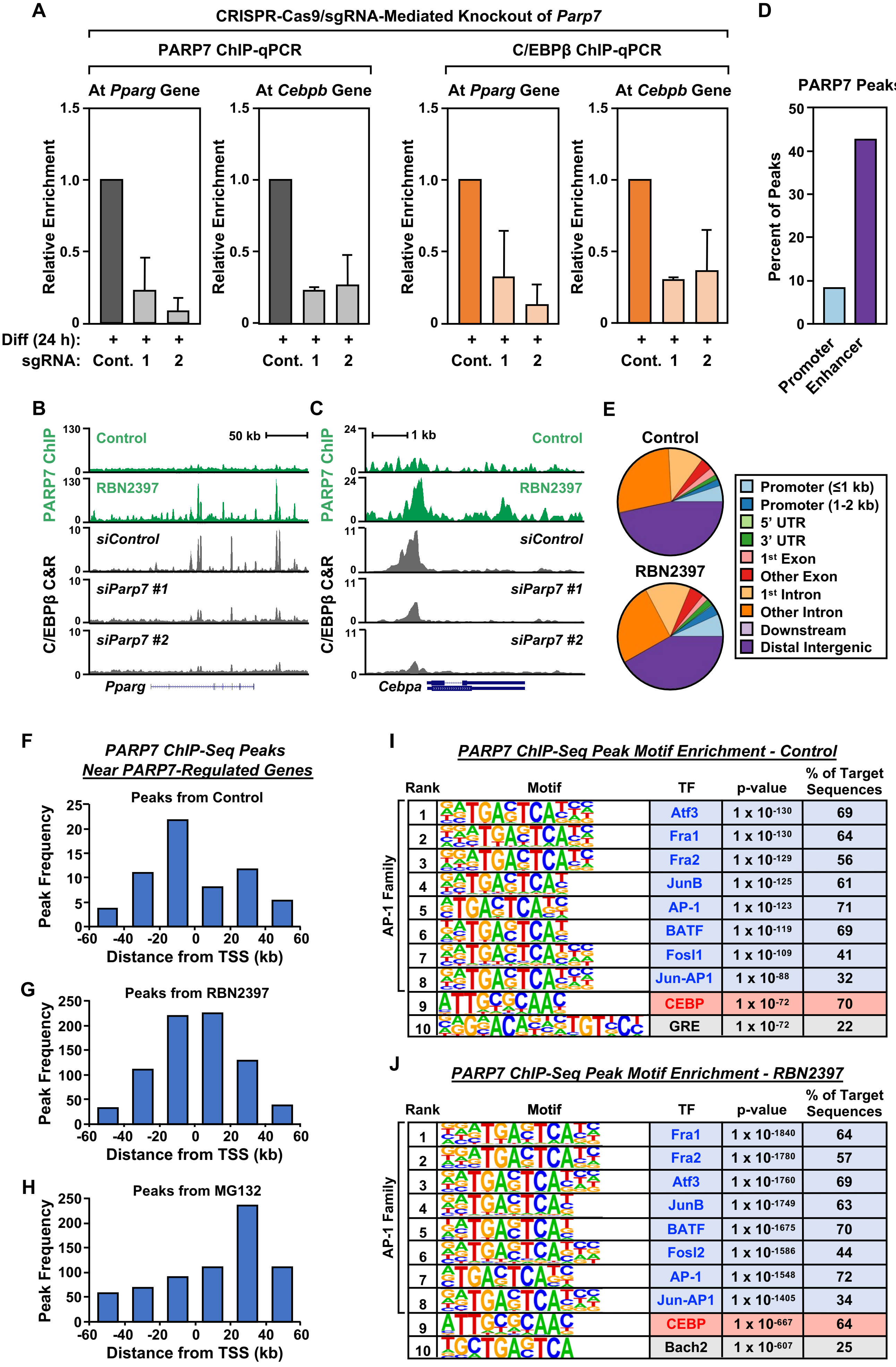


Suppl. Fig. S5 – Stokes *et al.* (2025)

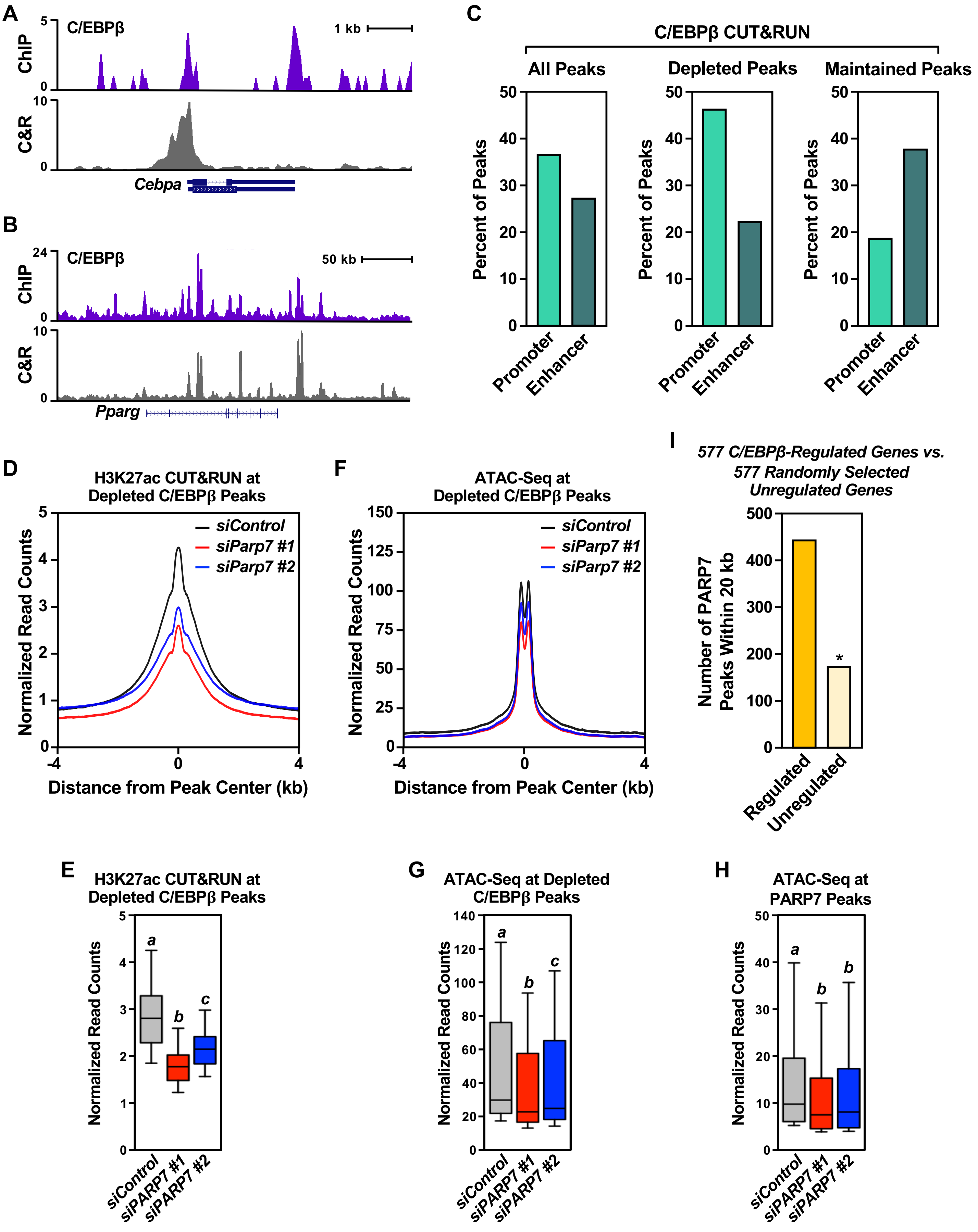


Suppl. Fig. S6 – Stokes *et al.* (2025)

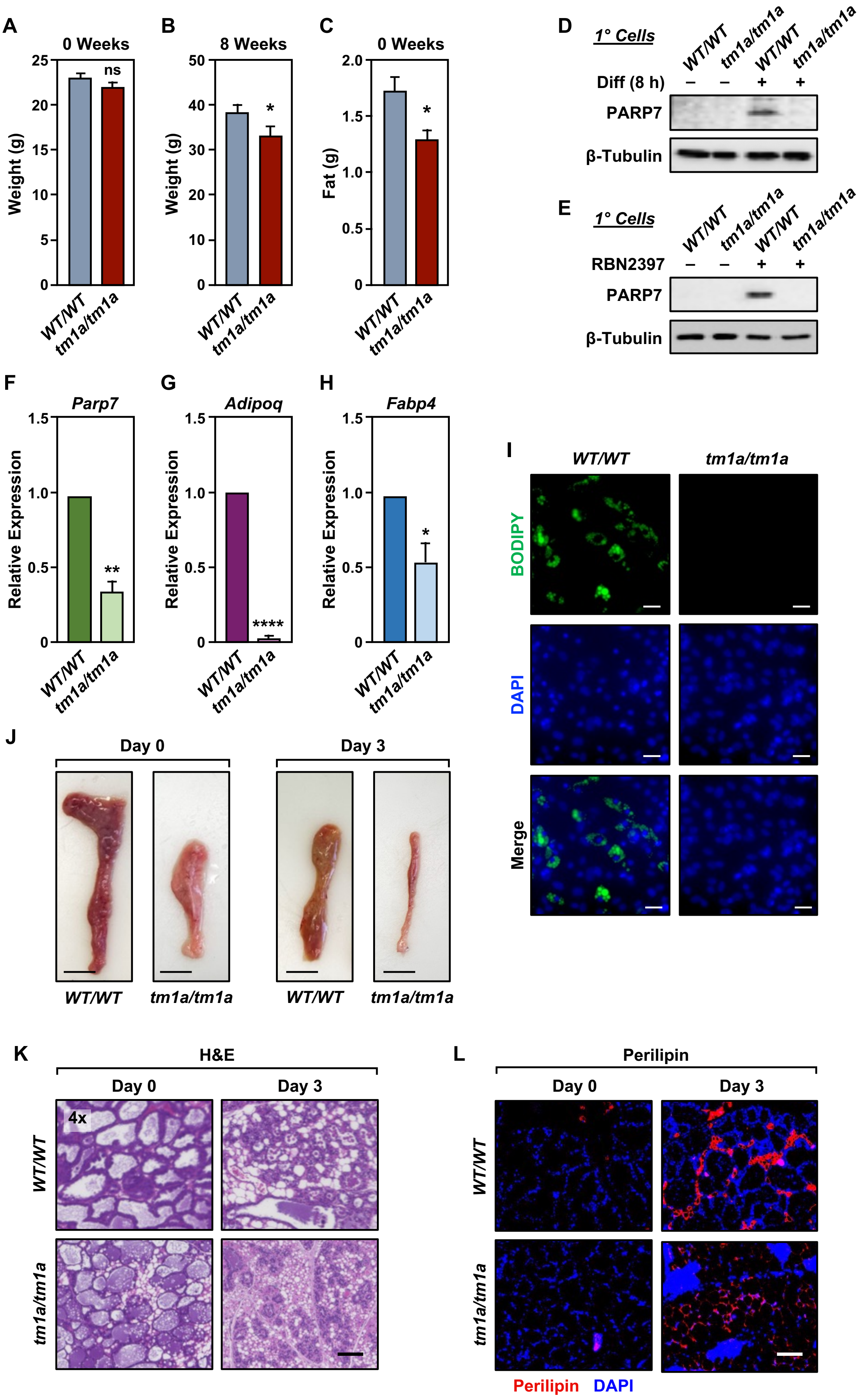


Suppl. Fig. S7 – Stokes *et al.* (2025)

**A**

| | Number of Litters | Litter Size | Percent Male | Percent Female | Percent *WT/WT* | Percent *WT/tm1a* | Percent *tm1a/tm1a* |
|---|---|---|---|---|---|---|---|
| Expected | n.a. | n.a. | 50 | 50 | 25 | 50 | 25 |
| *WT/tm1a x WT/tm1a* | 27 | 6.1 ± 0.5 | 50.8 ± 0.4 | 49.1 ± 0.2 | 35.3 ± 0.2 | 55.0 ± 0.4 | 10.7 ± 0.2 |
| C57BL/6 | 16 | 7.5 ± 0.7 | 54.1 ± 0.3 | 45.8 ± 0.5 | 100 | n.a. | n.a. |

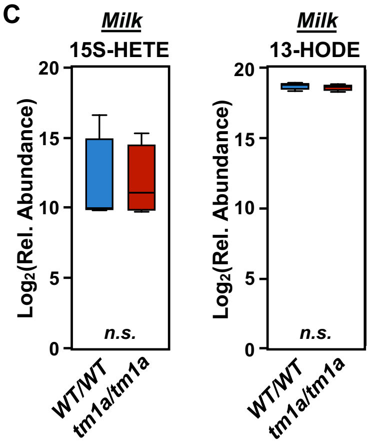


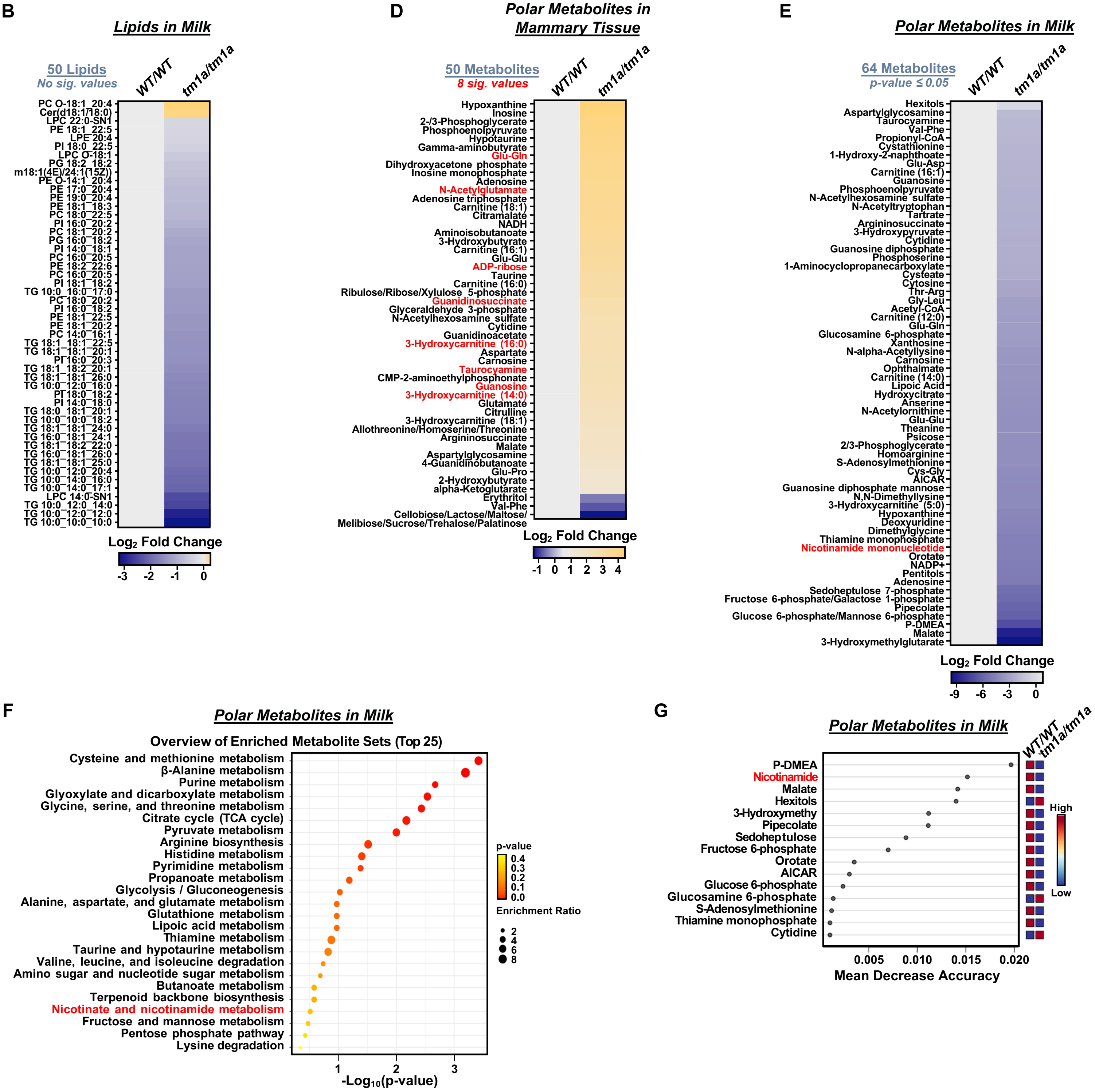


**Suppl. Fig. S8 – Stokes *et al.* (2025)**
